# Functional single cell selection and annotated profiling of dynamically changing cancer cells

**DOI:** 10.1101/2021.10.12.464054

**Authors:** Li You, Pin-Rui Su, Max Betjes, Reza Ghadiri Rad, Cecile Beerens, Eva van Oosten, Felix Leufkens, Paulina Gasecka, Mauro Muraro, Ruud van Tol, Ting-Chun Chou, Debby van Steenderen, Shazia Farooq, Jose Angelito U. Hardillo, Robert Baatenburg de Jong, Daan Brinks, Miao-Ping Chien

## Abstract

A method connecting single cell genomic or transcriptomic profiles to functional cellular characteristics, in particular time-varying phenotypic changes, would be transformative for single cell and cancer biology. Here, we present fSCS: functional single cell selection. This technology combines a custom-built ultrawide field-of-view optical screening microscope, fast automated image analysis and a new photolabeling method, phototagging, using a newly synthesized visible-light-photoactivatable dye. Using fSCS, we screen, selectively photolabel and isolate cells of interest from large heterogeneous populations based on functional dynamics like fast migration, morphological variation, small molecule uptake or cell division. We combined fSCS with single cell RNA sequencing for functionally annotated transcriptomic profiling of fast migrating and spindle-shaped MCF10A cells with or without TGFβ induction. We identified critical genes and pathways driving aggressive migration as well as mesenchymal-like morphology that could not be detected with state-of-the-art single cell transcriptomic analysis. fSCS provides a crucial upstream selection paradigm for single cell sequencing independent of biomarkers, allows enrichment of rare cells and can facilitate the identification and understanding of molecular mechanisms underlying functional phenotypes.

---

Tumor heterogeneity is a leading cause of failing cancer treatment^1, 2^. State-of-the-art single cell sequencing methods allow profiling whole genomes or transcriptomes of individual cells^3^. A technology that allows enriching and profiling rare and sparse subsets of cancer cells in tumor samples, that differentiates populations with mildly differing gene expression profiles and that allows linking of aberrant phenotypes to genomic or transcriptomic profiles^4, 5^ would have a transformative impact on cancer biology, as therapy resistant cells initially often exist in small quantities, gradually diverge in gene expression, show aberrant behavior (including e.g. aggressive migration^6, 7^ and morphological deviations^8^), and lack reliable biomarkers^9, 10^. Profiling these functionally diverse cells will allow unraveling the exact molecular mechanisms of distinct cancer driving behaviors of tumor subpopulations^1–3^.

We developed a functional single-cell selection pipeline (fSCS) to select well-defined subpopulations of target cells based on time-varying phenotypic changes, like fast migration or cell division, while maintaining cell viability. We profile those cells and correlate the sequencing information with functional characteristics (functional single cell sequencing, FUNseq). fSCS is used to identify and selectively label individual cells of interest (N=1 to 10^4^ simultaneously), from large heterogeneous populations of cells.

We created a high-throughput screening microscope, (Ultrawide Field-of-view Optical microscope, UFO, **Fig. S1**), which combines a large field of view of up to 42 mm^2^ with high spatial and temporal resolution (up to 0.8 μm/pixel, up to 30 ms/frame). Cells can be imaged in large quantities (~20k cells, **Figs. 1a, S2**) in both white light and fluorescence. We created a fast automated image analysis pipeline, capable of processing tens of thousands of cells simultaneously in fluorescence measurements of labeled cell nuclei (modified Tracking Gaussian Mixture model (mTGMM); **Figs. 1b**, **S3, Supplement**). The coordinates of cells of interest are fed to a digital micromirror device (DMD) or a pair of galvanometer scanning mirrors (galvo mirrors), which pattern light to selectively illuminate target cells. We achieved labeling of target cells with patterned light (**Fig. 1c**) using either genetically engineered photoactivatable (i.e., PA-GFP^11^) or photoconvertible proteins^12^ (i.e., mMaple3^13^), or photoactivatable dyes. To make fSCS suitable for expeditious single cell selection from biopsies or patient-derived primary cultures, and compatible with ubiquitous GFP reporter lines, we created a modified rhodamine-based photoactivatable dye. We conjugated a photosensitizer moiety, thioxanthene^14^, to the nitrophenyl-protected rhodamine (**Fig. S4**). We increased the photoactivation efficacy ~2.4 fold using visible 405 nm excitation^14^.

**Figure 1.**
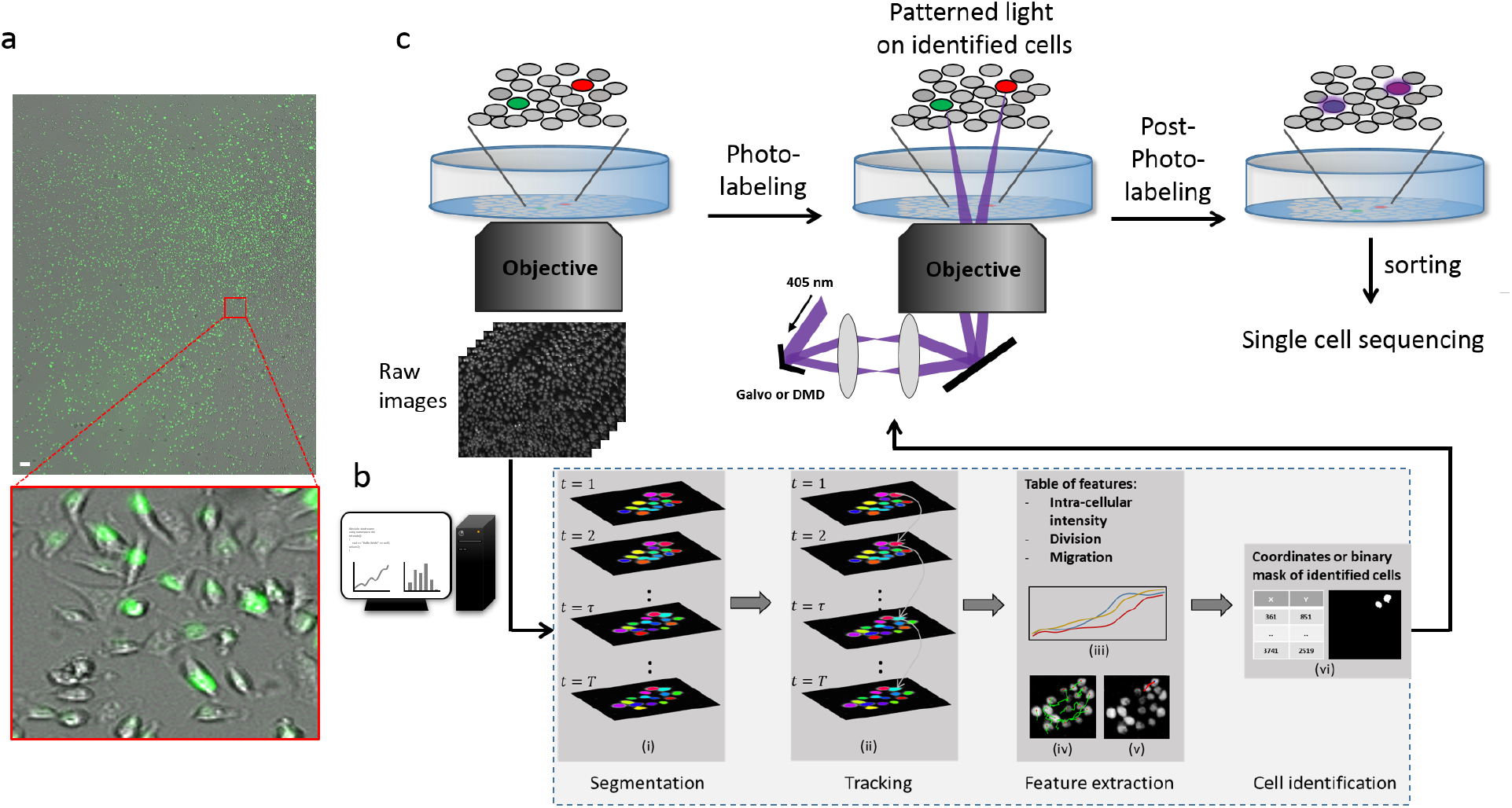
Functional Single Cell selection (fSCS) consists of ultrawide field-of-view miscroscopy, fast and accurate cell tracking, and target cell identification and photoselection. **a)** White-light and fluorescent image of MCF10A cells expressing H2B-GFP, imaged with NA 0.25. Image contains ~2×10^4^ cells. Scale bar 100 μm. Magnification is digital only, i.e. all information in the magnified image is contained in the parent image. **b)** Schematic of the software pipeline. (i) Images are pre-processed and segmented. (ii) Cell tracking is performed. (iii-v) From the post-tracking matrix cellular features are extracted including intracellular intensity changes (iii), cell migration (iv) and cell division (v). (vi) Coordinates of cells of interest are extracted. **c)** Schematic of the hardware pipeline. Movies are acquired on UFO and processed in real time using the mTGMM pipeline. Coordinates of cells of interest are extracted and sent to DMD or galvo mirrors for selective illumination, which locally converts the photolabeling reporter or dye. The photolabeled cells are separated via a cell sorter for downstream experiments like single cell sequencing.

We used CW lasers with either the DMD or galvo mirrors for phototagging in 2D cultures using 1-photon excitation. The DMD is particularly useful to label cells of interest in large quantities (**Fig. S5**, ~500 cells labeled in parallel). We used 100 fs pulses from a Titanium-Sapphire laser (Ti:Sapph) with the galvo mirrors for cell labeling in three-dimensional (3D) samples using 2-photon excitation (**Fig. S6**). We then separated the selectively labelled cells using a standard fluorescence-activated cell sorting machine for downstream studies like single cell sequencing.

The full fscs pipeline is illustrated in **Fig. 1.** We achieve a high tracking accuracy of 92.3 % (number of successful tracks per number of total tracks; **Supplement, Fig. S3**) and high sensitivity (97.6 %) and specificity (99.6 %) for cell division detection (**Supplement, Fig. S7**). We selectively label cells of interest with sensitivity of 97.4% and specificity of 99.9% (**Supplement, Fig. S8**). The activated phototagging dye is retained in target cells for a minimum of 12 hours in its photoactivated state (**Supplement, Fig. S9**). The photoselected cells maintain viability after the fSCS protocol (**Supplement, Fig. S10**) and can be re-cultured. We did not find any meaningful differentially expressed genes in randomly phototagged cells in comparison to non-phototagged cells in bulk cell RNA sequencing data (**Supplement, Fig. S11**), indicating that the phototagging procedure did not measurably impact the cells.

fSCS allows using any microscopically observable phenomenon, labeled or unlabeled, static or dynamic, as the criterion for cell selection. This includes intracellular dynamics (e.g. small molecule uptake, **Fig. 2a**) and cellular dynamics (e.g. division, **Fig. 2b**). To demonstrate genotypephenotype linking using FUNseq, we designed experiments targeted to discovering the genetic basis of fast migration and mesenchymal-like morphology transition in cancerous cells, as we hypothesized that these would be at the basis of metastatic behavior^6–8, 15^.

**Figure 2.**
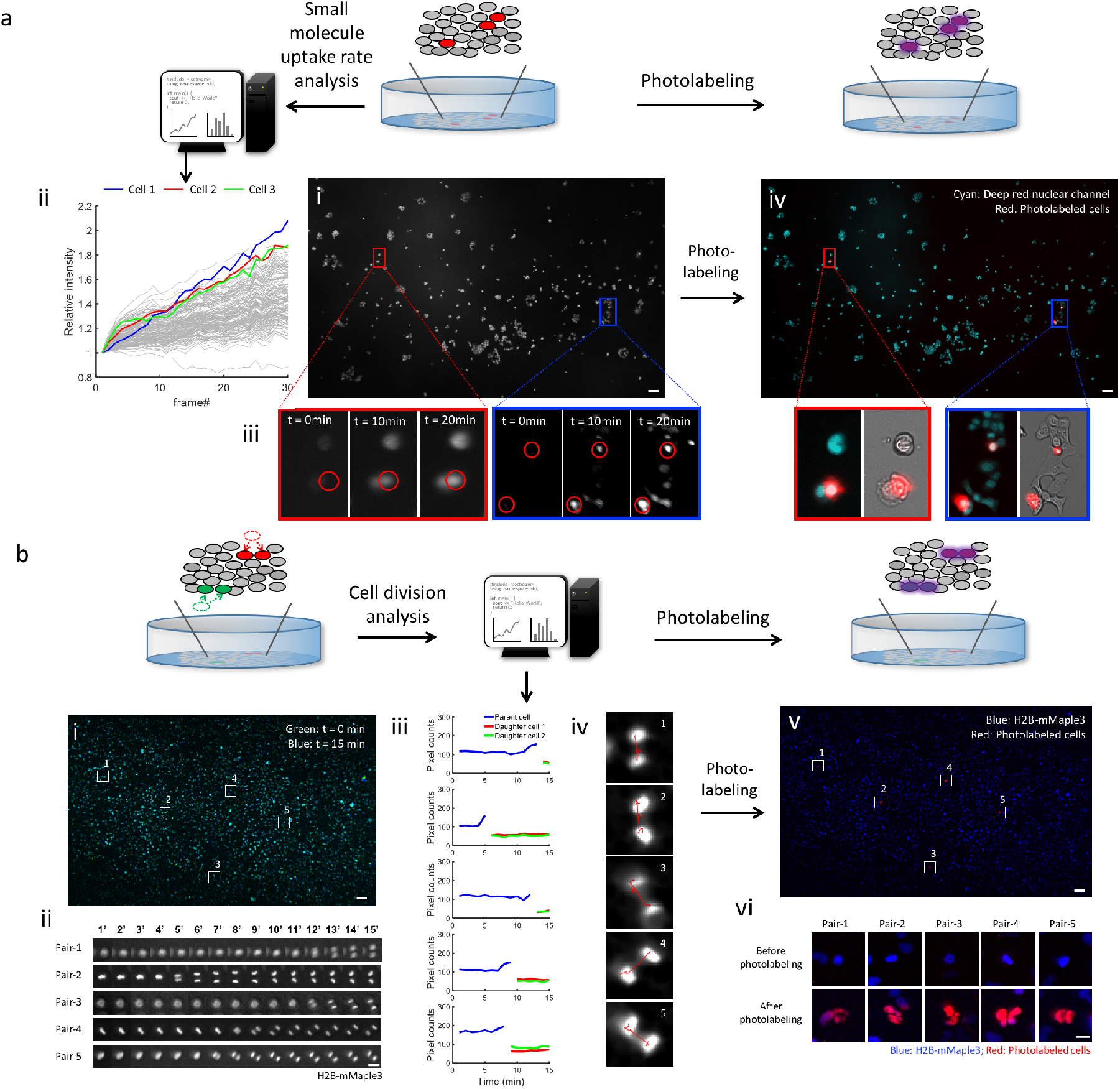
fSCS allows cell selection based on dynamically changing phenotypes. **a)** Cell selection based on small molecule uptake rate. a-i) Deep red nuclear dye was added to MCF10A-H2B-GFP cells. The sample was sequentially imaged using 460 nm and 637 nm excitation for 20 minutes at 6 frames/minute. a-ii) Dye uptake was tracked and the three cells with the fastest small molecule uptake rate were determined (highlight). a-iii) Magnified images of Cells 1-3. a-iv) Cells 1-3 were phototagged. **b)** Cell selection based on division rate. U2OS-H2B-mMaple3 cells were imaged using 460nm excitation for 15 min at 1 frame/minute. b-i) overlay of the first frame (green) and last frame (blue) of the movie. b-ii) 5 pairs of newly dividing cells were identified. b-iii) Size traces of 5 pairs of newly dividing cells; blue: parent cells; green, red: daughter cells. b-iv) Cell lineage trace of the newly divided cells. b-v) Newly divided cell pairs photolabelled (red) after photoconversion of mMaple3 with 405nm excitation. b-vi) Digitally magnified images of 5 pairs of newly divided photolabeled cells. Scale bars: b-i,v: 200 μm, b-ii, vi: 20 μm.

From a population of MCF10A-H2B-GFP cells, we monitored cellular migration trajectories (**Fig. 3a,b)** and detected cells with mesenchymal-like morphology (spindle shape; **Fig.3c-e**; **Supplement, Fig. S12**). We identified and selected the fast-moving cells **(Fig. 3b, Fig. S13)** and spindle cells **(Fig. 3e)**. We also induced the same cell line with TGFβ, an inducer of the epithelial-to-mesenchymal transition (EMT)^8^ in the hope of enhancing heterogeneity in both phenotypes and expression profiles^8^.

**Figure 3.**
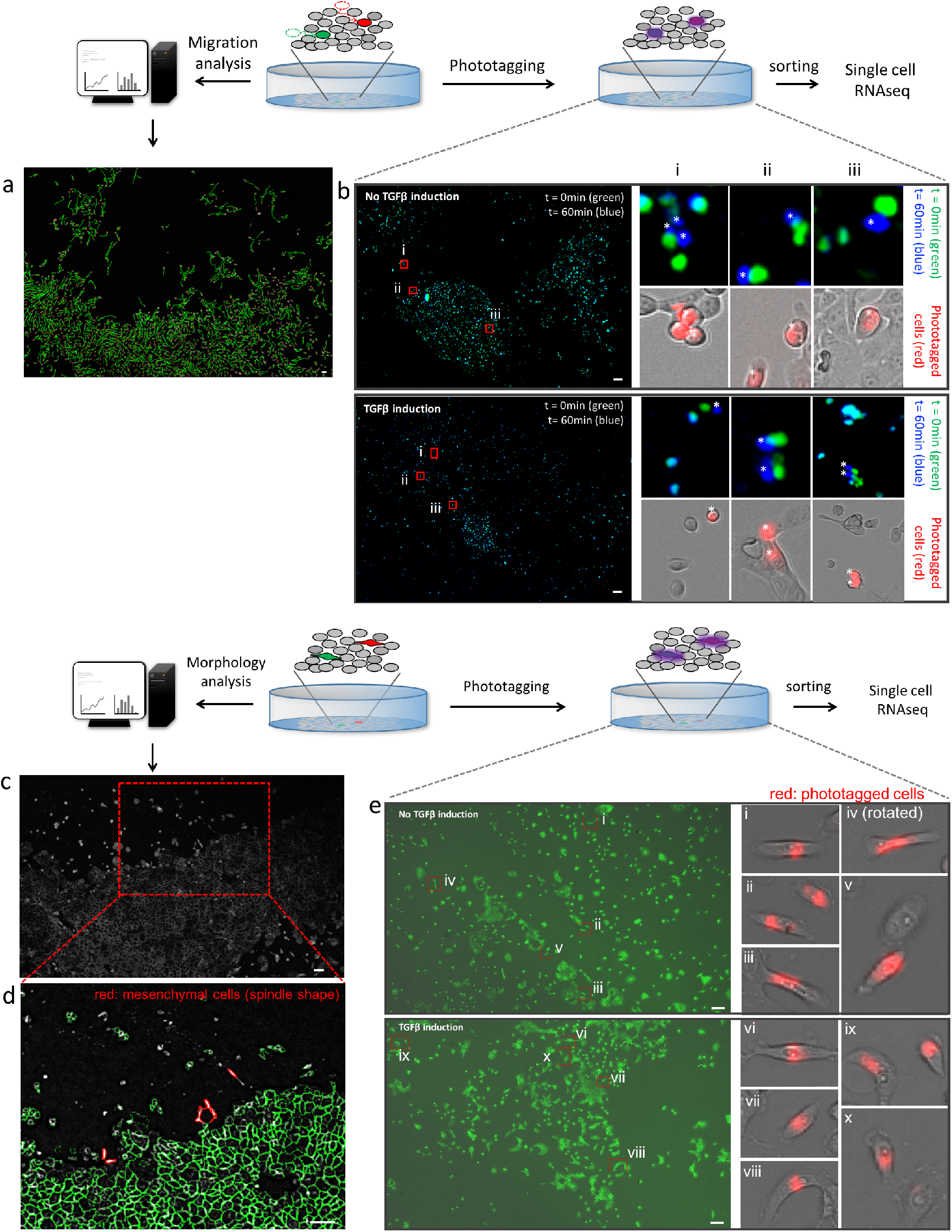
Identification, isolation and selection of MCF10A-H2B-GFP cells with fast migration and mesenchymal-like morphology. **a)** Migration trajectory of MCF10A-H2B-GFP cells analyzed and visualized by mTGMM. **b)** Overlay of position of non-TGFβ and TGFβ induced MCF10A-H2B-GFP cells at t=0 (green) and t=60 minutes (blue). Insets i-iii: six (non-TGFβ induced) and five (TGFβ induced) representative fast moving cells identified (indicated by *) after mTGMM and labelled via phototagging. **c)** MCF10A-H2B-GFP cells stained with Deep-red membrane dye. **d)** Image from c) analyzed and visualized after mTGMM allowing quantification of cellular morphology. **e)** Overlay of bright-field image and deep-red membrane stain image (green color) of non-TGFβ and TGFβ induced MCF10A-H2B-GFP cells. Insets i-v: five representative non-TGFβ induced cells with mesenchymal-like morphology identified after mTGMM and labelled via phototagging (red). Insets vi-x: five representative TGFβ induced cells with mesenchymal-like morphology identified after mTGMM and labelled via phototagging (red). Scale bars: 200 μm.

In a screen for migration speed we phototagged ~150-200 fast-moving cells (**Fig. 3b, Fig. S13**) from populations of ~20k cells. We then separated the phototagged cells using a fluorescence activated cell sorter (FACS) for subsequent single cell transcriptomic sequencing^16^ together with ~250 slow-moving cells. In a separate screen we phototagged and separated ~150-200 spindle-shaped cells (**Fig. 3e**) for analysis with ~ 250 non-spindle shaped cells.

Standard dimensionality reduction methods failed to identify any clear clusters associated with phenotypes of interest (**Fig. S14**). This is a known problem in unsupervised clustering analysis involving samples with rare or sparse cells, or populations with gradually changing gene expression profiles^17^. Even in fSCS enriched data, no well-defined clusters are formed (both noninduced and induced cells) (**Fig. 4a-d, Fig. S15**). FUNseq solves the clustering problem through functional annotation. Using the prior information provided by fSCS, one can distinguish clusters with more or less tagged cells (**Fig. 4a-d)**. We scored EMT^18^ in fast (phototagged) and slow (non-phototagged) moving cells and spindle (phototagged) and non-spindle (non-phototagged) cells, in both non-induced and induced conditions, based on an EMT hallmark gene set^19^ (**Fig. 4e,f**). We observe enrichment for tagged cells at high EMT scores (**Fig. 4e,f**). Supervised EMT analysis shows high coherence of EMT-marker expression and a consistent higher expression of these markers in cell groups with a higher proportion of tagged cells in all conditions (**Supplement; Figs. S16-S17**). The overlap between the two populations (tagged vs untagged) suggests that the failing clustering without the use of prior information is due to the gradual change in gene expression between the phototagged and non-phototagged cells.

**Figure 4.**
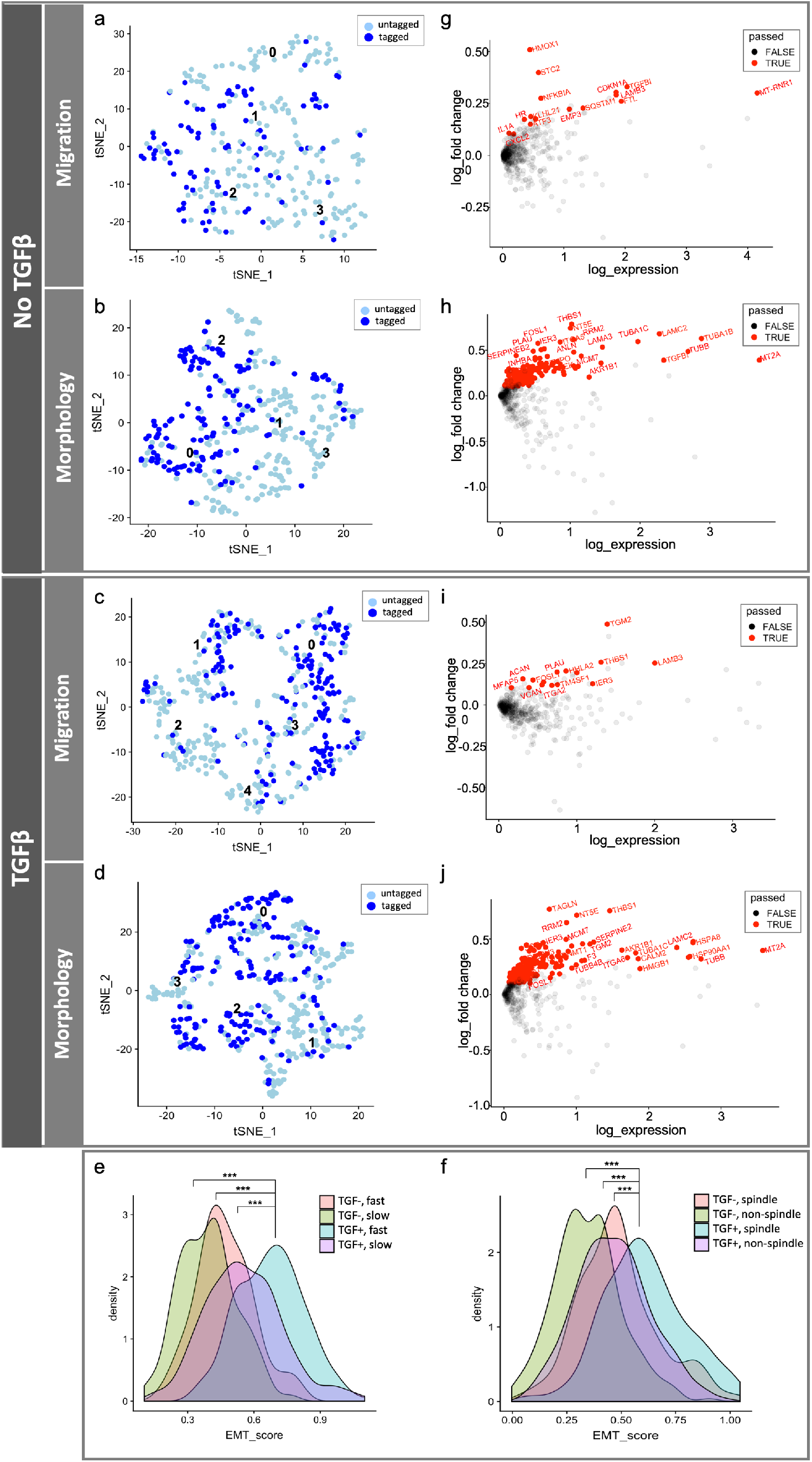
Functionally annotated single cell transcriptomic profiling (FUNseq) of MCF10A-H2B-GFP cells with fast migration and mesenchymal-like morphology. **a, b)** tSNE plot of non-TGFβ induced fast migrating (a) and spindle (b) cells. Dark blue: cells with fast migration (a) or mesenchymal-like morphology (b) (tagged); light blue: cells with slow migration (a) or endothelial morphology (b) (untagged). Numbers 0-3 (a,b) indicate the SNN cluster numbers. **c, d)** tSNE plots of TGFβ induced fast migrating (c) or spindle (d) and cells. Dark blue: cells with fast migration (c) or mesenchymal-like morphology (d) (tagged). light blue: cells with slow migration (c) or endothelial morphology (d) (untagged). Number 0-4 (c) or 0-3 (d) indicate the SNN cluster numbers. **e)** EMT score of fast (tagged) and slow (untagged) cells with or without TGFβ induction. EMT scoring against the EMT reference list from the EMT hallmark gene set shown in Subramanian et. al.^19^ Significance (*) was computed by Two-sample t-test (*** ≤ 0.001). **f)** EMT score of spindle (tagged) and non-spindle (untagged) cells with or without TGFβ induction. EMT scoring against the EMT reference list from the EMT hallmark gene set shown in Subramanian et. al.^19^ Significance (*) was computed by Two-sample t-test (*** ≤ 0.001). **g, h)** Smear plots of differentially expressed genes between tagged and untagged cells (**g,** migration data; **h**, morphology data) in the non-TGFβ induced condition. Upregulated genes that passed multiple testing correction (p<0.05) are shown in red. **i, j)** Smear plots of differentially expressed genes between tagged and untagged cells (**i**, migration data; **j**, morphology data) in the TGFβ induced condition. Upregulated genes that passed multiple testing correction (p<0.05) are shown in red.

Functional annotation allows direct analysis of differential gene expression between phototagged and non-phototagged cells, bypassing clustering methods completely. In non-TGFβ-induced, fast migrating cells, we identified two driving pathways: TGFβ and NF-κB (based on the overexpressed genes TGFBI and NFKBIA, ATF3, IL1A, CDKN1A, CXCL2, SQSTM1 and LAMB3 (hallmark gene set^19^); **Figs. 4g, S18a, S19a**). Both TGFβ and NF-κB were also identified as pathways driving spindle shape in non-TGFβ-induced conditions, based on the differentially expressed genes TGFBI and IER3, PLAU, FOSL1, INHBA and SERPINEB2 (hallmark gene set^19^; **Figs. 4g**, **S18a**, **S19b**). In spindle cells, differential gene expression analysis also identified Tubulin and Laminin structural proteins (TUBA1B, TUBA1C, TUBB, LAMC2 and LAMA3 genes), indicating these playing a critical role in maintaining spindle shape (**Fig. 4h**, **S19c**). Markedly, neither these genes nor TGFβ or NF-κB pathways were identified using clustering methods followed by differential gene analysis without functional annotation (**Fig. S15 i,j**).

In the TGFβ-induced condition, among TGFβ and NF-κB pathways, only the NF-κB pathway was identified in the fast-moving cells (IER3, LAMB3, PLAU and FOSL1 genes (hallmark gene set^19^), **Fig. 4i, S18b**, **S19d**). In the spindle cells, several NF-κB pathway associated genes, though not the full pathway, were identified as significantly overexpressed (IER3, F3 and FOSL1 genes (hallmark gene set^19^), **Fig. 4i**, supplementary methods and **Figs. S18b, S19e**). As all cells were induced with TGFβ, it is expected that TGFβ genes did not show up in a differential expression analysis. Tubulin and Laminin structural proteins were again identified in spindle cells (TUBA1C, TUBB, TUBB4B and LAMC2; **Fig. 4j**, **S19f**). Once again, these genes and pathways were not identified through clustering methods followed by differential gene analysis (**Fig. S15 k,l**).

Validation of the role of TGFβ and NF-κB pathways in fast migration and mesenchymal-like morphology confirmed that fSCS correctly linked phenotype to driving genotype. We conducted a gene knockdown (RNA interference) assay on MCF10A cells (**Supplementary methods**) and found a strong decrease of fast migrating and spindle shaped cells in populations with knocked down TGFβ and NF-κB (**Fig. S20**).

TGFβ- and NF-κB-mediated pathways are among those that have been reported to drive EMT^8, 20^ (including among others WNT, Notch, HIF and Ras-ERK). EMT describes a collective biological process that can drive tumor aggressiveness and lead to various cancer-driving phenotypes, but the specific link between individual pathways and phenotypes often remains unclear^8, 20^. The above findings indicate that specifically TGFβ and NF-κB pathways dominantly drive tumorigenic fast migration and transition to mesenchymal-like morphology even in non-TGFβ induced conditions. Moreover, only through FUNseq, we directly link the main driving pathways (TGFβ and NF-κB) to the observed aggressive phenotypes (fast migration and mesenchymal-like morphology) in individual cell populations, using limited sample sizes for the assay and direct readout. In this diagnostically important context, these pathways were hidden without fSCS annotation-assisted analysis.

By combining ultrawide field-of-view microscopy with real-time accurate cell tracking and phototagging, fSCS and FUNseq makes it possible to study the intra-population heterogeneity of phenotype-driving pathways in (patient-derived) tumor samples and thereby opens the possibility for targeted and personalized interventions.

fSCS allows enrichment of rare cells and functional annotation, which lets us identify TGFβ and NF-κB as pathways that cause fast migration and mesenchymal-like morphology independent of external TGFβ induction in MCF10A cells. Most importantly, fSCS and FUNseq allows linking genotypes with any phenotypes of interest observable under a microscope. The method allows profiling dynamically aggressive cells, however sparse, such as cancer stem-like cells or cancer-driving cells, for which we lack reliable or universal biomarkers^9, 10^, facilitating the understanding of cancer-driving and therapeutic mechanisms.

## METHODS

### Ultrawide Field-of-view Optical Microscope (UFO) setup

UFO is a custom-built microscope, which incorporates a large chip-size CMOS Point Grey camera (GS3-U3-123S6M-C, 4096×3000 pixels, 3.45 μm/pixel, FLIR) and large field-of-view (FOV) objectives (Olympus MVP Plan Apochromat, 1x and 0.63x) (**Fig. S1**) with comparatively high numerical aperture (for the 1x objective, NA= 0.5; for the 0.63x objective, NA=0.25).

This microscope images a 7.3 × 5.7 or 3.65 x 2.83 mm FOV with 1.7 μm or 0.8 μm spatial resolution, respectively, at 30 ms temporal resolution, which is sufficient for recording most cellular dynamics. Fast temporal dynamics can be reached with a downsized FOV.

Illumination is provided by several CW laser lines (405nm (2W, MDL-HD-405, CNI), 460nm (800 mW, MDL-III-460, CNI), 532nm (1500mW, MGL-FN-532, CNI) and 637nm (1.3W, MDL-MD-637, CNI)). These lasers can be projected to samples with a custom-designed 45° AOI collimated, 3.3 x 2.2” trichroic mirror (R_avg_: 405-460nm, 785-1300nm and R_abs_: 532/637 nm; Alluxa). Fluorescence was filtered through a custom-designed 2” tri-band emission filter (Od6_avg_: 400-465nm / 527-537nm / 632-642nm / 785-1300nm; Alluxa) and collected with an MVX-TLU tube lens (Olympus Telan lens). The custom-made trichroic and emission filters are designed in such a way that switching filters between experiments is not required. All illumination (except 405nm) is temporally structured by an acousto-optic tunable filter (Gooch & Housego) and all illumination is modulated spatially by a digital micromirror device (DMD, V-9501 VIS, Vialux). The DMD is reimaged onto the sample with targeted illumination at 4.8 μm spatial resolution and 0.1 ms temporal resolution.

For point-scanning illumination via mirror galvanometers (15mm clear aperture mirror, 6240H, Cambridge Technology), 200mW 405nm laser (MDL-XS-405, CNI) and Titanium-Sapphire 100fs pulsed laser (Coherent Mira900) were implemented and collimated at the back focal plane of the objective. The light was then defocused to obtain 10 μm spot at the sample and was steered in the sample plane using galvanized mirrors in a conjugate plane.

### Cell cultures

#### MCF10A-H2B-GFP

MCF10A-H2B-GFP breast epithelial cells, a gift from Dr. Reuven Agami (Dutch National Cancer Institute, NKI), were grown in DMEM/F-12 supplemented with 5% horse serum, 1% penicillin/streptomycin, EGF (10ng/mL), Hydrocortisone (500ng/mL), Cholera Toxin (100ng/mL) and insulin (10 μg/mL) in a 37°C incubator under 5% CO_2_. For TGFβ1 treatment, MCF10A cells were treated with human recombinant TGFβ1 (R&D Systems) for 2 weeks.

Before conducting experiments, 10,000-20,000 cells were seeded on fibronectin (0.1 mg/mL)-coated 10mm-glass bottom dishes in the MCF10A culture medium (described above) without phenol red. Experiments were performed 16-24 hours after plating on the glass-bottom dishes.

#### U2OS cell culture

U2OS-H2B-mMaple3 cells were grown in DMEM medium supplemented with 10% fetal bovine serum and 1% penicillin/streptomycin in a 37°C incubator under 5% CO_2_.

Before conducting experiments, 30,000 cells/cm^2^ were seeded on gelatin (0.1 mg/mL)-coated glass bottom dishes in the DMEM culture medium (10% fetal bovine serum, 1% penicillin/streptomycin) without phenol red. Experiments were performed 16-24 hours after plating on the glass-bottom dishes.

### mTGMM

Image analysis computation was performed on a single workstation, Dell Precision 7920, with the following hardware components: dual Intel(R) Xeon(R) Gold 6130 CPUs, 32GB DDR4-2666MHz memory, a Nvidia GeForce 1080 GPU.

mTGMM is based on the Tracking Gaussian Mixture model^21^. In the first step, raw images are preprocessed (for images with heterogeneous fluorescence intensity profile between cells: imtophat -> contrast stretching -> Gaussian smoothing; for images with relatively homogeneous fluorescence intensity profile between cells: imtophat->Gaussian smoothing).

Every image t ∈ {1,2, …,T] is processed in parallel for nuclei segmentation, using a modified watershed approach. We record centroids and masks for individual cell-nuclei at t. Foreground pixels are grouped into superpixels (i.e., one superpixel is a set of connected pixels that does not overlap any other superpixel). The superpixels are trimmed using local Otsu’ thresholding, to determine the relation between superpixels and nuclei: superpixels that are still connected will be grouped into one nucleus, while an isolated superpixel represents a nucleus itself. In other words, a nucleus may have one or multiple superpixels. We implemented a parameter to adjust a threshold value (in number of pixels) of contacted pixels between two Gaussian Mixture Models (GMMs, indicating two nuclei). This value is particularly sensitive to cell division and should be tuned accordingly in different sample types.

The intensity profile of a nucleus is modelled as a 2D Gaussian distribution. Nuclei tracking is done by forwarding every Gaussian from time point *t* to *t* + 1 using Bayesian inference, with a priori knowledge that the position, shape, overall intensity of nuclei cannot change dramatically between two consecutive time points. Based on the tracking information we generate a feature table with records of cellular migration, division and intra-cellular intensity (details in **Supplement**).

### Synthesis of phototagging dye

The two starting materials, photoactivatable rhodamine (PA-Rho; *orřho*-nitroveratryl-oxycarbonyl-5-carboxy-Q-rhodamine) (Sigma) and 2-amino-thioxanthone (Key Organics), were combined in a one-step synthesis where the thioxanthone amine coupled to the carboxylic group of PA-Rho to form an amide conjugation (**Supplement, Fig. S4**). We added 2-amino-thioxanthone (17.2 μmol, 2 equ) in dimethylsulfoxide (DMSO, 0.1 mL, Sigma) to a solution of PA-Rho (8.6 μmol, 1 equ) in DMSO (0.8 mL) with 1.5 equ of HATU coupling reagent (in DMSO, 0.1 mL; Sigma) and 4 equ of diisopropylethylamine (Sigma). After stirring for 16 hrs under nitrogen, the product was separated from unreacted starting materials via Semi-Prep HPLC (retention time: 23.1 min; RP-C18 column with a 30 min gradient of 50% to 100% ACN containing 0.1% TFA). The product is confirmed via LC-MS (Exp. m/z: 1142.84, [M+H^+^]: 1142.11). The product fluorescesces around 580 nm which makes it suitable for use with GFP reporter lines, unlike common photoactivatable and photoconvertible proteins.

### Photolabeling experiment

#### with phototagging agent

Cells of interest, determined through mTGMM, were selectively photolabeled via phototagging or photoconvertible mMaple3. For phototagging, cells pre-incubated with 15 μM of phototagging reagent (diluted in no phenol-red culture medium) were selectively photolabeled using 405nm light (100 J/cm^2^).

Upon selective illumination, the phototagging compound inside of the target cells will be photoactivated and can be visualized by 532nm excitation (25 mW/cm^2^) after photoactivation (activated phototagging dye: lex: 532nm, lem: 553nm). Cells without 405nm activation will remain dark upon 532nm excitation.

#### with photoconvertible mMaple3

For photolabeling using photoconvertible mMaple3, U2OS cells expressing H2B-mMaple3 were used. Before photoconversion by 405nm (10 J/cm^2^), mMaple3 can only be imaged with blue light (460nm in the UFO setup, 10 mW/cm^2^). Upon photoconversion by 405nm, mMaple3 can be additionally excited and visualized by 532nm excitation (15 mW/cm^2^). Cells without 405nm photoconversion will remain dark upon green light excitation.

### Single cell RNA sequencing analysis

Single cells were collected in two 384 well-plates (three repeats for the migration data; two repeats for the morphology data). After sequencing and QC we had data from 465 TGFβ induced cells for the migration assay and 431 TGFβ induced cells for the morphology assay. For the experiment on non-TGFβ induced cells, 302 cells were analyzed in the migration assay and 375 cells in the morphology assay, for a total dataset of 1573 deep sequenced cells (~100k reads/cell). SORT-seq sequencing and read alignment were performed as described by Muraro *et al*^16^ (Single Cell Discoveries).

#### Quality control

All the single cell analysis is performed in Seurat v3^22, 23^. The logarithm of the feature count formed a clear bimodal distribution (**Fig. S21a,c**). Cells with the feature counts larger than 2,500 and fewer than 10,000 were selected. In addition, cells with a percentage of mitochondrial counts lower than 30% were selected, again selecting one part of a bimodal distribution (**Fig. S21b,d**).

#### Processing and feature selection

The analysis was performed on the TGFβ-induced and non-TGFβ-induced cells separately. Counts were log normalized using Seurat v3’s *NormalizeData* function followed by batch effect correction. Batch effects were removed using Seurat v3’s canonical correlation analysis and anchor cell identification method (the *FindIntegrationAnchors* function followed by *IntegrateData* function) ^22, 23^, after which variance associated with the replicate was removed. Feature selection was then performed using the *FindVariableFeatures* function (variance-stabilizing transformation method) from Seurat v3^23^. A subset of 3000 variant genes were selected for the following analysis.

#### Cell cycle regression

For the cell cycle scoring, a set of S and a set of G2/M-phase markers are used^18^ and the S.Score and G2M.Score were computed using the *CellCycleScoring* function in Seurat v3. For cell cycle regression, cell cycle genes were regressed out using the *ScaleData* function *(vars.to.regress = c(“S.Score”, “G2M.Score”))* in Seurat v3.

#### Dimensionality reduction and clustering

After cell cycle regression, we performed Principle component analysis (PCA) on the subset of highly dispersed genes. We chose the first 5 principle components for SNN-clustering based on the jackstraw procedure^24^ (**Fig. S22**). Both in the induced and non-induced data, we saw a loss of significance in PC’s after the fifth principal component (**Fig. S22**).

SNN-clustering was performed with the *FindClusters* function in Seurat v3^22, 23^. The resolution parameter was swept in the range of 0.6 to 1.4 (**Fig. S23 (migration), S24 (morphology)**) (0.6 to 1.2 should typically yield good results according to the package authors). For all other parameters defaults were used.

#### Module scoring

For the EMT scoring we used a module of the EMT hallmark gene set described in Subramanian et. al.^19^. The scoring was done with the *AddModuleScore* in Seurat^22, 23^. For a given cell, this compares the expression of every gene in the module with similar genes (based on average expression in the whole set).

#### Differential gene expression analysis

Differentially expressed genes between the tagged (fast or spindle cells) and untagged (slow or non-spindle cells) groups (functional annotation) were accessed through the *FindMarkers* function in Seurat v3^22, 23^ using the *Model-based Analysis of Single-cell Transcriptomics* (MAST) method^25^ (Bonferroni adjusted P value < 0.05).

## Supporting information

FUNseq_Supplementary information

## ACKNOWLEDGEMENTS

MPC acknowledges support from the Oncode Institute, Cancer GenomiCs.nl (CGC), NWO (the Netherlands Organization for Scientific Research) Veni Grant, and Erasmus MC grant. DB acknowledges support by an NWO Start-up Grant (740.018.018) and ERC Starting Grant (850818 - MULTIVIsion). We thank Reuven Agami and Maarten Paul for the kind gift of MCF10A-H2B-GFP and U2OS-H2B-mMaple3 cell lines, respectively. We thank Arjan Theil and Tsung Wai for technical assistance with FACS sorting. We thank Jaco Kraan and John Martens for the support of using their cell separation machine. We thank Erasmus MC Center for Biomics for the bulk-cell transcriptomic sequencing. We thank KT Chen for the phototagging purification. We thank Philipp Keller for discussions about TGMM.

## AUTHOR CONTRIBUTIONS

LY scripted the mTGMM cell tracking algorithm and analyzed most of the image analysis data. MB scripted and analyzed the single cell sequencing data. PRS designed the experiment and scripted the algorithm for intracellular dynamics measurement. PG conducted the 3D phototagging experiment. CB contributed to all the cell culture preparation for the experiments and the gene knock-down experiments. EvO, FL and TCC contributed to the morphology detection algorithm. MM advised on some of the single cell experiment and sequencing analysis. DvS conducted part of the cell migration and sorting experiment. MPC and DB set up data-acquisition hardware and software with contributions from RvT, SF and PG. MPC, DB, LY, MB and PRS wrote the paper with input from all authors. MPC and DB initiated, contributed to and supervised all aspects of the project.

The data that support these findings are available from the corresponding author upon reasonable request.

The algorithms used in this manuscript will be available on Github.

