## Supplementary material for "Functional single cell selection and annotated profiling of dynamically changing cancer cells": FUNseq_Supplementary information

### - Supplementary methods
### - Figures S1-S26
### - Supplemental references

### Detailed mTGMM analysis procedure

UFO allows high-throughput cell screening to record time-varying functional cellular dynamics (i.e., fast migration or cell division) as well as intracellular dynamics (i.e., small-molecule uptake rate). A fast, automated and accurate cell tracking algorithm is required to process these dynamics in an efficient manner. In samples of migrating cells, postprocessing or an insufficiently fast algorithm would lead to targeting coordinates from which the targets cell has moved away already. For example, cells with a size of ~40 μm can partly migrate away (i.e., with speed of ~1 μm/min) from their original locations after 20min of movement. Therefore, any cellular or intracellular dynamics analysis should be completely done within this timeframe. To achieve this, we modified one of the fastest cell tracking algorithms, a Tracking Gaussian Mixture Model (TGMM, Ref. 10 in the main text), and made it suitable for use with 2D samples (**Fig. S3**). We named our algorithm, modified TGMM (mTGMM).

Raw images acquired from UFO are immediately sent to pre-processing, segmentation, tracking, feature extraction and cell identification (**Fig. 1b**, main text).

In the first step, raw images are pre-processed (for images with heterogeneous fluorescence intensity profile between cells: imtophat -> contrast stretching -> Gaussian smoothing; for images with relatively homogeneous fluorescence intensity profile between cells: imtophat->Gaussian smoothing). Heterogeneous gene expression shown in different fluorescence intensity between cell to cell is a known phenomenon (**Fig. S25**) and often caused imperfect cell segmentation due to inhomogeneous fluorescence intensity. Instead of a median filter used in TGMM, we first used imtophat to remove inhomogeneous illumination profile followed by a contrast-stretching intensity transform shown in Eq. 1, where $I_{new}$ is a newly transformed image from $I_{original}$, the original image. The image is thresholded at the intensity m and the steepness of the threshold is given by S.

$$I_{new} = 1./(1+(m./(double(I_{orignal})+eps)).\text{^}S \quad \text{(Eq. 1)}$$

The newly adjusted image is then smoothed by a Gaussian filter (with a standard deviation at 0.5 or 1) to slightly increase fluorescence blurriness, which can improve Gaussian model fitting used in the subsequent analysis.

The original TGMM offered an option to run a 2D sample by creating a fake 3D sample via inserting a black slice below a 2D image and used the 3D connectivity (6, 26, 74) for processing this fake 3D sample. In our modified mTGMM, we used a similar approach to create a so called pseudo-3D sample: Let $f(x, y)$ be the intensity of a 2D image $t \in \{1, \dots, T\}$ at coordinates $(x, y) \in \mathbb{Z}^2$, $g(x, y, z)$ be the intensity of the pseudo-3D image $t$ at $(x, y, z) \in \mathbb{Z}^3$, then we have

$$g(x, y, z) = \begin{cases} f(x, y) & if\ z = 0, \\ 0 & if\ z = -1. \end{cases}$$

We then applied 4-connected or 8-connected pixel neighborhood in this pseudo-3D sample. We also offer this variable in the software configuration file for users to adjust.

Every image $t \in \{1, \ldots, T\}$ will be in parallel processed for nuclei segmentation, including individual cell-nucleus' centroid and mask at t. A modified watershed approach is used for fore-ground detection, and these foreground pixels which consist of an object (i.e., one nucleus or a part of it) will be grouped into superpixels (i.e., one superpixel is a set of connected pixels that does not at all overlay every other superpixel). The superpixels will be trimmed using the local Otsu' thresholding, as shrinking the segmentation mask, so to determine the relation between superpixels and nuclei: superpixels that are still connected will be grouped into one nucleus, while an isolated superpixel represents a nucleus itself. In other words, a nucleus may have one or multiple superpixels. We implemented a parameter, called *minNeighboringVoxel,* in mTGMM's configuration file to adjust a threshold value (in number of pixels) of contacted pixels between two nuclei. This value is particularly sensitive to cell division and should be tuned accordingly in different sample types.

The intensity profile of a nucleus will be modelled as a Gaussian distribution in 2D. The assignment of a Gaussian per cell nucleus will be followed by the step of tracking. Every Gaussian will be forwarded from time point $t$ to the next $t + 1$ using Bayesian inference, with a priori knowledge that the position, shape, overall intensity of nuclei cannot change dramatically within two consecutive time points. In other words, a cell-nucleus' segments at each $t \in \{1, \ldots, T\}$ are temporally sequenced. A feature profile (i.e., parameters) per cell nucleus will be built, including its 2D centroid, 2D covariance matrix (shape), its id, its superpixel id and its parent id.

Next, the feature tables, which respectively record cells' migration, division and intracellular intensity information, will be generated based on the tracking output. To examine migration, we extract distance-based features such as *Net Distance, Accumulated Distance, Maxi Distance* (between two consecutive time points) that a cell travelled, and/or orientation-based features such as *Turning angle*. To study division, we follow the lineage and record the time and position when a cell splits into two daughter cells. Cell nucleus size dynamics shall follow a typical trend before/when/after it divides into two, and thus in our pipeline conditions are imposed to exclude false positives caused by under- and/or over- segmentation. The mTGMM pipeline outputs an index matrix $A = (a_1, \ldots, a_N)^{\mathrm{T}} \in \mathbb{Z}^{N \times T}$for retrieving cell lineage, where $N$ is the cell count at the last time point, $T$ is the time duration of imaging, and $a_i$ (row vector) denotes the $i$-th cell's index over all $t \in \{1, \ldots, T\}$. Two daughter cells inherit their parent's indices when the parent disappears right after a division. To accurately detect the newly dividing cells, we implemented a few restraints in mTGMM to ensure a high accuracy of detection, one of which is the size of the parent cell must be bigger than both daughter cells, and the size of both daughter cells must be similar.
In **Fig. 2b** in the main text, we report division detection using our method. To follow intracellular dynamics, we obtain the mask per nucleus per time point, and select the cells that have undergone a dramatic intracellular change. We present this experiment is **Fig. 2a** in the main text. mTGMM offers the visualization of segmentation and tracked trajectory, which is critical to examine the performance of the analysis (**Fig. S3**).

Last, our cells of interest will be selected according to the sorted feature tables. These are the cells either with fast migration (i.e., which move further than a pre-determined distance threshold), newly divided, or fast small molecule uptake rate (i.e., whose intracellular intensity changes greater than a pre-defined grayscale value). The coordinates of these target cells will be passed to the microscope. The galvo mirrors or DMD can then steer light selectively to these target cells.

The whole mTGMM pipeline is illustrated in **Fig. 1b.** Time-lapse raw images are acquired and immediately processed through the computational workflow. Steps from left to right (i-vi): (i) parallel cell-nucleus segmentation, i.e., every image at $t \in \{1, \dots, T\}$ is simultaneously processed for nuclei segmentation, including individual nuclei's centroid and mask (i.e., pixels that consists of nuclei) at $t$. (ii) sequential linkage of cell-nucleus over time, i.e., every individual cell-nucleus is followed, building a link among a cell-nucleus' segments at each $t \in \{1, \dots, T\}$ . (iii-v) migration, division and intracellular intensity profiles are extracted from the output of the tracking matrix, i.e., three tables record cell-nuclei's intracellular dynamics (iii), migratory trajectory (iv) and division (iv), respectively. For fast computation, one may skip the visualization part. (vi) Fast moving cells (i.e., their migratory distance is larger than a pre-determined threshold), spindle cells, proliferative cells, or fast small molecule uptake cells (i.e., their intracellular intensity changes greater than a pre-defined threshold) are identified as cells of interest. Those cells' coordinates are extracted and sent to UFO's galvo mirrors or DMD for selectively photolabeling of cells of interest (**Fig. 1c**).

### Mesenchymal-like morphology detection

Both nuclei-stained (H2B-GFP) and membrane-stained images (CellMask Deep Red plasma membrane stain; 1x working solution, ThermoFisher) were used to detect the spindle MCF10A-H2B-GFP cells. Nuclei were imaged by blue light (460nm in UFO, 10 mW/cm$^2$) and the membrane stain was imaged by red light (637 nm in UFO, 15 mW/cm$^2$).

Our method to extract morphology features (per cell membrane) consists of three steps:

*First*, nuclei detection is described above using nuclei-stained images via mTGMM.

*Second*, cell membrane is detected using max-flow algorithm[1]: conceptually speaking, looking for the object edge that cuts an image into foreground (e.g., bright pixels) and background (e.g., dark pixels) equals to finding the cut that gives the maximum amount of flow (i.e., difference of grayscale values between adjacent pixels) of that image. Initially some pixels are labelled as either foreground or background (i.e., in our case, we make use of the detected nuclei and label their pixels as foreground for subsequent computation). Then the algorithm goes through every unlabeled pixel and its neighboring ones, and takes the best guess on the label of this pixel. The best guess is the result of maximizing a flow function[2] which can achieve a globally optimal result. The max-flow algorithm is sensitive to image quality (such as uneven illumination, high noise-to-signal ratio, blurred edges, etc), hence beforehand membraned-stain preprocessing (i.e., top-hat filter to remove background illumination, ridge enhancement to smooth and enhance edges, etc) is required.

*Thirdly*, morphological properties of cell membranes are quantified, and these properties include: (1) circularity, i.e., a ratio equal to $4\pi A/C^2$ where $A$ is the area of a membrane and $C$ is its perimeter. (2) solidity, i.e., a ratio between the area of the convex hull and the membrane area. Usually, epithelial cells are convex and this ratio is 1. Spindle cells can be concave with a solidity smaller than 1. Please note when mesenchymal cells are convex (i.e., ellipse-like shape), this property will not play a role. (3) eccentricity: an ellipsoidal representation of a membrane area is computed using the matrix of inertia. The eccentricity is defined as the ratio between the focal distance and the length of the major axis of the ellipse, ranging from 0 for a circle to 1 for a line. For an epithelial cell this value approximates 0 whereas for a mesenchymal cell this value rises as a result of elongation. Eccentricity turned out to be dominant factor distinguishing spindle from non-spindle cells. Spindle cells were defined as cells with eccentricity=>0.95.

### Sensitivity and Specificity of mTGMM tracking algorithm

For quantification, SirDNA-stained MCF10A cells were used and cellular migration and division were monitored for 5 hours, generating a sequence of 151 frames (i.e., time interval 2 minutes) **Fig. S3 and S7**). This image sequence was immediately sent to mTGMM. To quantify the performance of mTGMM, we answer the following questions:

For an individual cell, we say its tracking is successful if it is tracked continuously over time whenever it persists. Precisely, we give the definition of accurate tracking of an individual cell: Let $\mathcal{P}(i,t) \in \mathbb{R}^2$ be the ground-truth position of cell $i$ at time $t$, $p(i,t) \in \mathbb{R}^2$ be the detected position of cell $i$ at time $t$. The tracking of cell $i$ is accurate if $p(i,t) = \mathcal{P}(i,t)$ for each $t \in \{t_0, \dots, T\}$, where $t_0$ denotes the first time point a cell appears and $T$ denotes the last time point of imaging. Thereby, the tracking accuracy is defined as $\frac{\#.\ successful\ tracks}{\#.\ total\ tracks}$.

mTGMM reaches 92.3 % tracking accuracy. A visualization of the migratory trajectory per cell is given in **Fig. S3**.

For the particular field-of-view (511 cells; **Fig. S7**), there were 41 pairs of newly dividing cells, of which 40 pairs were accurately detected and 2 other pairs were falsely detected (True positives: 40, True Negatives: 468, False Positives: 2, False Negatives: 1). This yields a high sensitivity (97.6 %) and high specificity (99.6 %) (**Fig. S7**). mTGMM allows users to manually delete the falsely detected cells from the analysis result.

For spindle cell detection, MCF10A-H2B-GFP cells stained with the CellMask Deep Red plasma membrane stain were used. For the particular field-of-view (2259 cells; **Fig. S12**), there were 22 cells showing mesenchymal-like morphology (spindle shape), of which 19 cells were accurately detected and 1 cell was falsely detected. (True positives: 19, True Negatives: 2239, False Positives: 1, False Negatives: 3) This yields a decent sensitivity (86.4 %) and high specificity (99.9 %) (**Fig. S12**). mTGMM allows users to manually delete the falsely detected cells from the analysis result.

### Computational efficiency of mTGMM tracking algorithm

Using our workstation and mTGMM, the average computational time for an image sequence with 4096 ×3000×30 (i.e., pixels x number of frames) and around 8500 cells per frame was 41 seconds for pre-processing, 106 seconds for segmentation, 102 seconds for tracking, and 20 or 113 (= 20

+ 93) seconds for feature extraction (20 seconds for calculating inter-cellular features such as migration and proliferative features, and another 20+93 seconds for extracting intra-cellular features based on segmentation masks).

mTGMM processes around 57,000 cells per minute on average to select fast-moving, proliferative or spindle cells, and the processing includes pre-processing, segmenting, tracking and cells-of-interest identifying. To target the intercellular dynamics which takes extra time, mTGMM processes around 42,000 cells per minute on average. Please note that, if the number of pixels per frame, and/or number of cells per frame decreases (e.g., a small field of view image with 512 ×512 pixels and about 100 cells), the computational efficiency will considerably increase. Throughout this paper, we report the computational time based on our large FOV samples.

Compared to commercialized software CellProfiler, which processes (segments and tracks) around 4,100 cells per minute, mTGMM is significantly faster.

**Phenotype persistence study (fast migration and mesenchymal-like morphology)**

To ensure that the phenotypes (fast migration and mesenchymal-like morphology) are persistent rather than stochastic events, we measured the correlation of the phenotype between the early time points and later time points (no phototagging). We observed that migration speed at early time points (t = 1 hr) highly correlates with migration speed at later time points (t = 3 hr and 6 hr, r = 0.85 and 0.73), and that 92.5% of cells with mesenchymal-like morphology maintain their spindle shape after 6 hr (**Fig. S26**).

**Photoactivation efficacy comparison between phototagging dye and the original photoactivatable-rhodamine dye**

To compare the photoactivation efficacy between phototagging dye and the original photoactivatable-rhodamine dye (PA-Rho), MCF10A cells incubated with 15 μM of both dyes. Using the same excitation energy (100 J/cm$^2$) via 405nm laser, phototagging dye (imaged by 532nm excitation, (25 mW/cm$^2$)) reaches a 2.4-fold higher fluorescence intensity than the PA-Rho dye, indicating 2.4 times more efficient photoactivation.

**Phototagging sensitivity and specificity**

To measure the phototagging sensitivity and specificity, we co-cultured a sparse amount (~2%) of MCF10A-H2B-GFP with MCF10A cells without GFP expression. We then selectively phototagged the cells expressing H2B-GFP to calculate the sensitivity and specificity of phototagging procedure. We identified 117 MCF10A-H2B-GFP cells in a population of ~ 5000 cells, of which 114 cells were successfully phototagged (see a representative image in **Fig. S8**). In addition, 8 cells were falsely phototagged, resulting in high sensitivity (97.4%) and high specificity (99.9%) of the phototagging method.

**Phototagging in 3D tumorsphere**

A tumorsphere was prepared in a 3D microwell plate (GravityTRAP-ULA, Perkim Elmer). ~3000 cells of MCF10A and MCF10A-H2B-GFP cells (500:1) were mixed and grew in the plate for a week before imaging. 15 uM of phototagging dye was pre-incubated with the tumorsphere for 20 min in the phenol-red free medium and rinsed away before the experiment. In **Fig. S6**, 6 GFP-expressing MCF10A cells (green) were identified in one of the z-planes, we then selected 3 of the 6 cells to phototag. After the phototagging procedure, 3 cells were successfully photoactivated by two-photon pulsed laser (0.65W, Mira900, Coherent) and could be imaged by 532nm green light excitation (25 mW/cm$^2$), shown in red in **Fig. S6**, i-iii. The tumorsphere was then stained with Deep Red NuclearMask (1x working solution, ThermoFischer) and imaged by 637 nm excitation (15 mW/cm$^2$) for visualizing all the MCF10A cells in the tumorsphere (shown in blue in the **Fig. S6**).

**<u>Retention time of activated phototagging dye</u>**

To measure how long the activated phototagging dye stayed inside of target cells, we measured the fluorescence intensity of activated phototagging dye on 200 randomly phototagged MCF10A cells over time. Intensity was obtained by averaging photocounts measured by camera with exposure time 1 sec over the area of the cell and subtracting the background level in the same channel. The fluorescence intensity gradually decayed over the period of 12 hrs as shown in **Fig. S9**, indicating that the photolabeled signal can be used up to 12 hrs for downstream cell separation.

**<u>Cell viability after phototagging</u>**

To ensure cells maintain viability and their genetic profile are not influenced after phototagging, we have conducted three experiments.

First, the cell viability staining (Life Technologies, Part Number: L-3224) assay was conducted. 140 MCF10A cells were randomly phototagged, (**Fig. S10**; the insets show 10 cells zoomed in) After phototagging, the whole dish was stained with Live Cell stain (Calcien AM, Invitrogen), which can be excited by blue light (460nm in UFO, 10 mW/cm$^2$) and imaged at the green channel. Live cells can be successfully stained with the Calcein AM dye as shown in **Fig. S10** (shown in green). In this figure, the phototagged cells (red, imaged by 532nm excitation (25 mW/cm$^2$) maintained viability (green) after the procedure.

Lastly, we compared the whole transcriptome sequencing data to ensure the genetic expression profiles were not impacted by phototagging. The PCA plot in **Fig. S11a** shows that randomly phototagged cells and non-phototagged cells have similar genetic expression profiles as they grouped with each other. Most importantly, we did not find any differentially expressed genes that passed significance testing (adjusted p value < 0.05, **Fig. S11b,c**), indicating that the phototagging treatment is very mild to cells.

**<u>Bulk cell RNA-sequencing</u>**

Total RNA was extracted from four populations of cells: two populations were MCF10A-H2B-GFP cells with TGFβ induction treated with and without phototagging, the other two populations were non-TGFβ induced MCF10A-H2B-GFP cells treated with and without phototagging. Three repeats were performed in each condition (>~1 million cells per repeat per condition). Phototagging was induced on a per-dish basis by flood illumination of the entire dish. Total RNA from each population was extracted using a RNAeasy plus Micro RNA kit (Qiagen) according to manufacturer's recommendations followed by sequencing (Erasmus MC Biomics Center). Libraries of all samples were prepared with the Illumina TruSeq Stranded mRNA Library Prep Kit. The resulting DNA libraries were sequenced according to the Illumina TruSeq Rapid v2 protocol on an Illumina HiSeq2500 sequencer. Reads were generated of 50 base-pairs in length and mapped to the GRCh38 human reference sequence using HiSat2 (version 2.1.0). Gene expression values were called using htseq-count (version 0.9.1). The analysis of the bulk cell sequencing data was done with the DESeq2 package[3]. The design formula was specified using the phototagging (phototagged versus non-phototagged) and the experimental conditions (TGFβ induced versus non-induced) as independent covariates. Significance testing was done without prior filtering or any independent filtering of genes.

### Single cell functional selection based on cellular and intracellular dynamics

We designed an experiment targeting fast small-molecule uptake rate of human breast epithelial cells stably expressing GFP (MCF10A-H2B-GFP). Upon small molecule addition (Deep Red nuclear dye), cells were immediately recorded/imaged sequentially in green (555-615 nm fluorescence) and far-red channels (660-770 nm fluorescence) (**Fig. 2a, i**) for 20 minutes at a framerate of 6 frames/min.

The GFP signal (green channel) of the cells was used to determine coordinates and cell-classified pixels of each cell. Increased intensity in the far-red channel reflected the amount of small molecule dye taken up by the cell (**Fig. 2a, ii-iii).** The coordinates of the 3 cells with fastest small molecule uptake were retrieved and sent to the galvo mirrors for phototagging (**Fig. 2a, iv**).

To demonstrate selective identification and photolabeling of cells based on cellular dynamics (i.e. behavioral, phenotypic differences between cells), we show the ability to select cells based on cell division (**Fig 2b**). U2OS-H2B-mMaple3 cells were imaged using 460nm excitation for 15 min at 1 frame/minute; 5 pairs of newly dividing cells were identified. Newly divided cell pairs were photolabelled after photoconversion of mMaple3 with 405nm excitation.

### Supervised EMT analysis

Supervised EMT analysis was performed to assess coherence of the EMT-scoring module. EMT-scores were used to group cells in eight bins with equal amounts of cells. Inside these bins all raw gene UMI counts were summed and normalized by the total number of UMI's in the bin. These UMI frequencies per genes are then scaled by the standard deviations across all bins. We then plotted all members of the module of EMT-genes. Hierarchical clustering was done on the gene expression profiles in the same way as in McFaline-Figueroa, J.L. et al.[4]

### Gene set enrichment analysis (GSEA)

The gene set enrichment analysis was conducted by the *fgsea* algorithm implemented in R[5, 6]. GSEA calculates an enrichment score, which quantifies the relevance of the hallmark gene set[7] to a particular group of selected genes (differentially expressed genes between tagged and untagged cells). The normalized enrichment score was analyzed to account for unequal gene set sizes and differences in the correlation between the gene sets and the expressed genes. The p value was calculated for each gene set, followed by Bonferroni multiple hypothesis test, providing the reported adjusted p-values.

### Gene knockdown assay (RNA interference assay, Fig. S20)

For small interfering RNA (siRNA)-mediated knockdown of TGFβ or NFkB gene, MCF10A-H2B-GFP cells were transfected with 10 nM of either the targeting or control siRNA (SMART pool, Horizon) using Lipofectamine RNAiMAX (Invitrogen) for 24 hrs. After that, medium was replaced with regular culture medium (no phenol red) for another 48hrs before imaging (migration and morphology).
After imaging, protein expression after siRNA gene knockdown was validated using Western blot analysis (**Fig. S20c**): cells were harvested and disrupted using Laemnli buffer, protein concentration was determined with Lowry assay. SDS-PAGE was performed on a 12% gel and blotted onto PVDF membrane, incubated with primary antibodies (***anti-NFkB antibody***, Abcam ab32360; ***anti-TGFβ antibody***, Abcam ab179695, ***Anti-β-Catenin antibody*** (internal control), BD Transduction Laboratories 610153) and secondary antibodies conjugated with HRP (Jackson Imm. Res. 515-035-003). Blots were developed with ECL and imaged on a AI600 Chemiluminescent Imager.

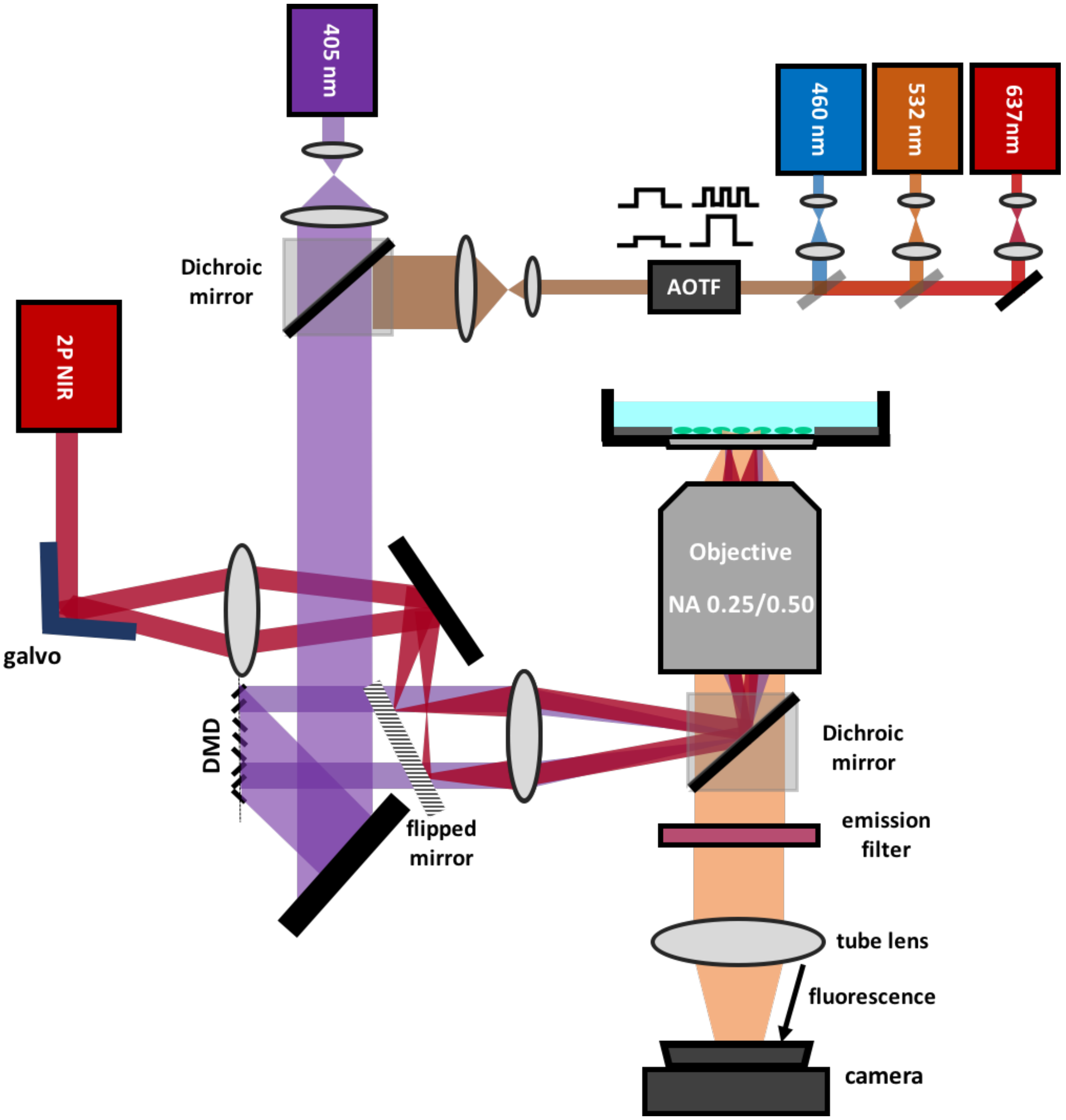


**Figure S1. A custom Ultrawide Field-of-view Optical Microscope (UFO microscope) for fSCS.** Large Field of view (FOV) objectives are combined with custom-made dielectric filters and a high pixel-density sCMOS camera. Spatial light patterning is achieved using a digital micromirror device (DMD) or galvanometer mirrors (galvo). Temporal structuring of illumination light at 460 nm, 532 nm and 637 nm is achieved using an acousto optic tunable filter (AOTF). The device combines a 405 nm CW laser for 2D patterning using 1-photon excitation and 100 fs, 810 nm pulses from a Titanium-Sapphire (Ti:Sapph) laser for 3D patterning using 2-photon excitation.

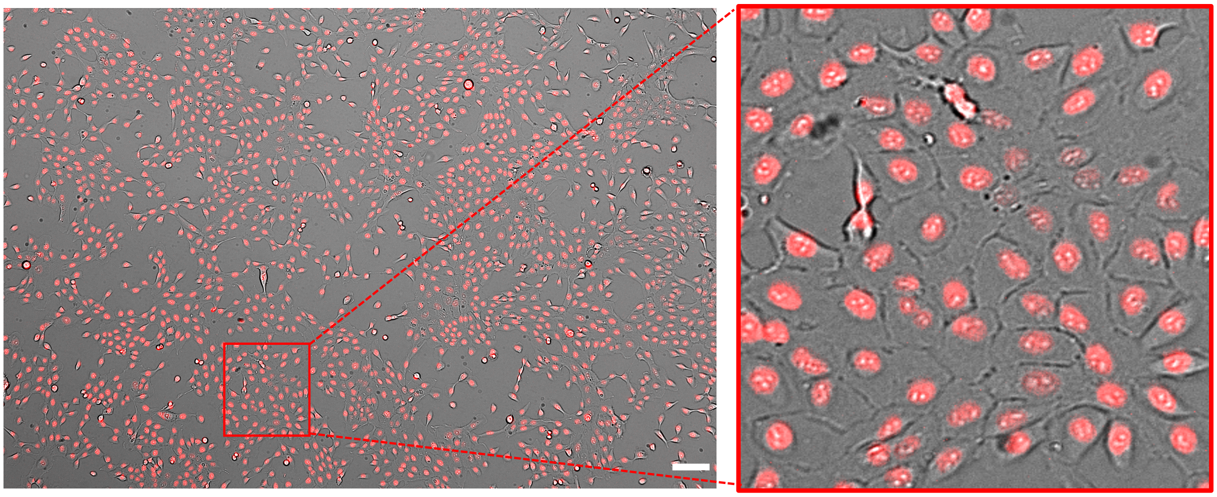

**Figure S2. High resolution, widefield imaging on UFO.** White-light and fluorescent images of SirDNA (0.5 $\mu$M)-stained MCF10A-H2B-GFP cells, imaged with NA 0.5. Image contains ~4,500 cells. Scale bar 100 µm. Magnification is digital only, i.e. all information in the magnified image is contained in the parent image.

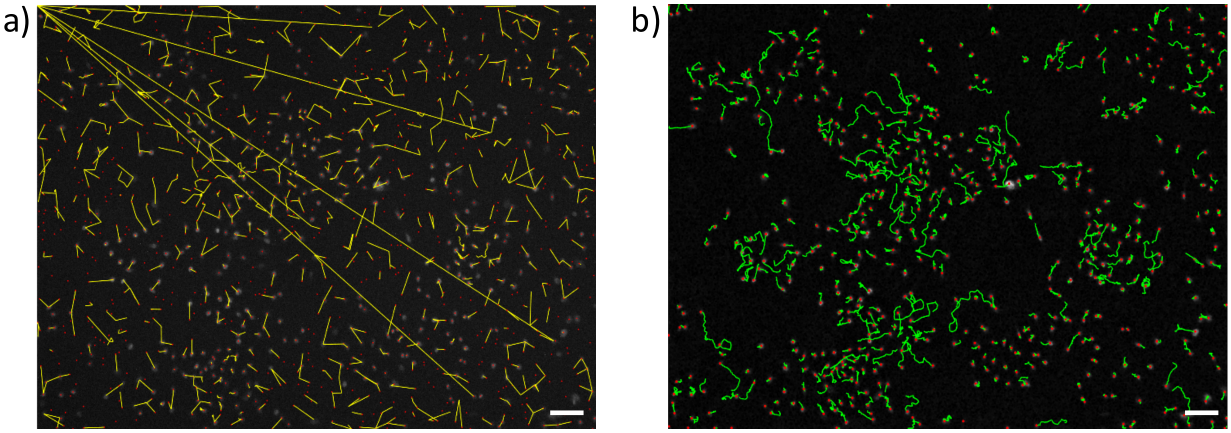


**Figure S3. Visualization of tracking (*Cell line: MCF10A + siRDNA stain*)**. a) Migratory trajectory (yellow) per cell (red) as output by the original TGMM, where over-segmentation and wrong links were observed. b) Migratory trajectory (green) per cell (red) as put out by mTGMM. In both cases we used the same image sequences. Scale bar 100 µm.

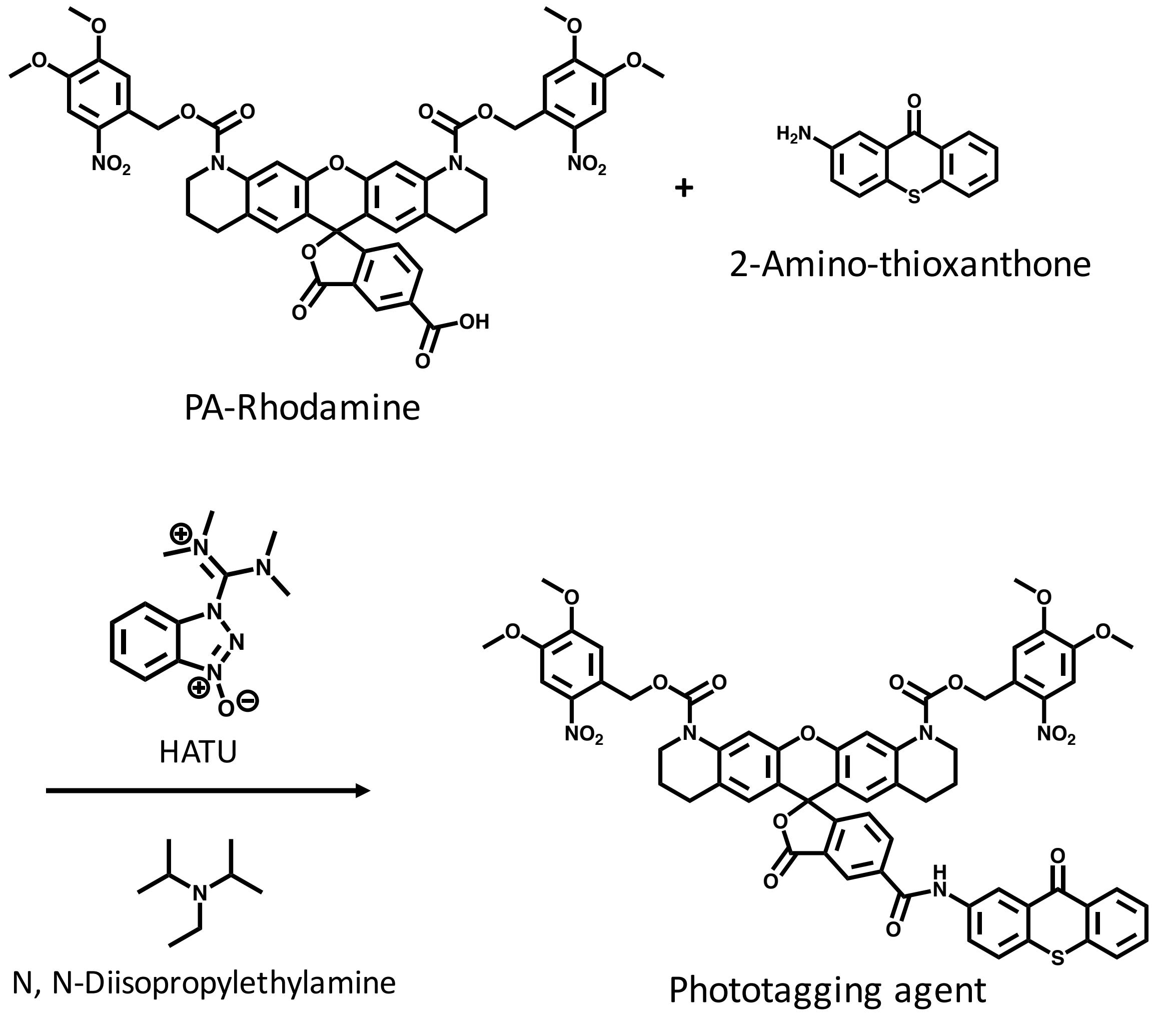


**Figure S4**. **Synthesis of phototagging agent**. Sulfo-Photoactivatable Rhodamine (PA-Rho, 1 equ) and 2-Amino-thioxanthone (2 equ) with HATU coupling agent (1.5 equ) and Diisopropylethylamine (4 equ) reacted in DMSO under nitrogen for 16 hrs. The product was purified by RP-C18 Semi-prep HPLC.

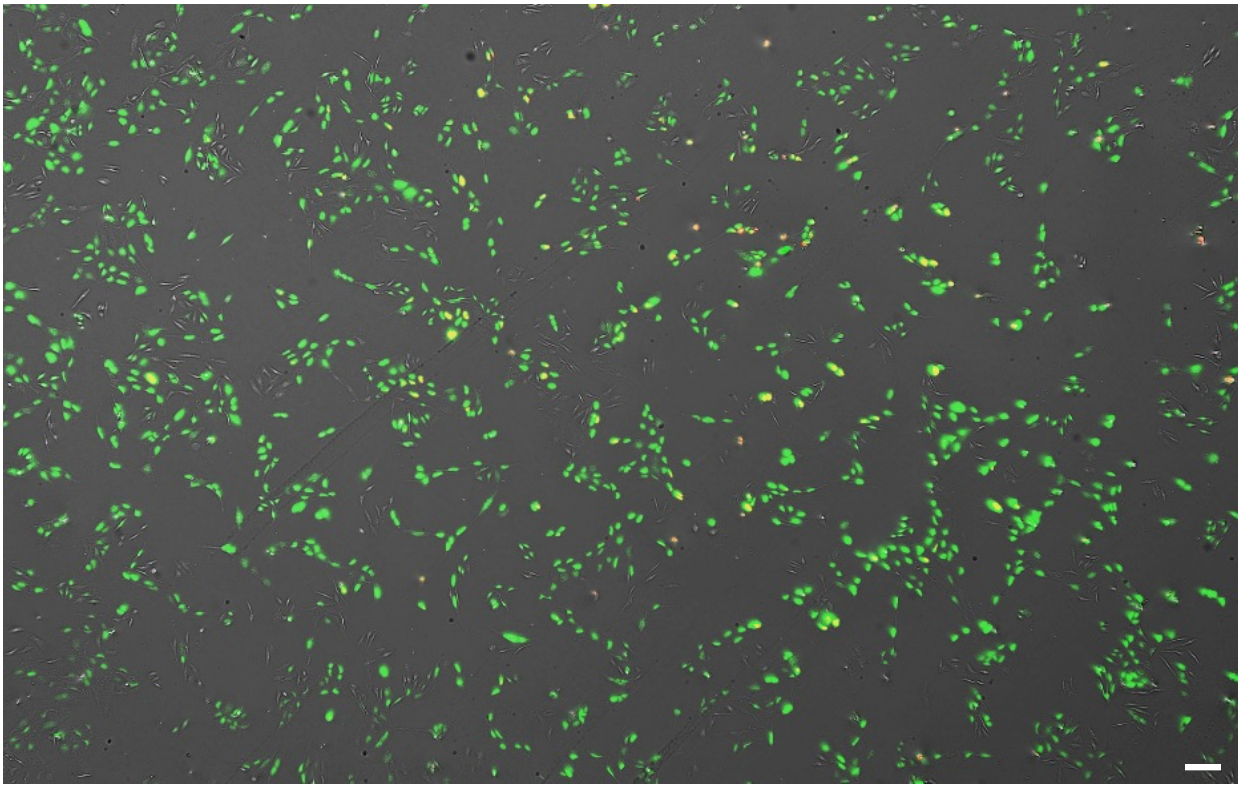

**Figure S5. Photolabeling of fast migrating U2OS-H2B-mMaple3 cells.** U2OS cells that migrated >10 $\mu$m (12 pixels; 0.8 $\mu$m/pixel) in an hour were identified after mTGMM (477 cells out of 1550 cells in this field-of-view) and successfully photolabelled (red) after 405nm (10 J/cm$^2$) activation using DMD. Green: U2OS cells. Red: Photoconverted U2OS cells (imaged by 532nm excitation). Scale bar 100 µm.

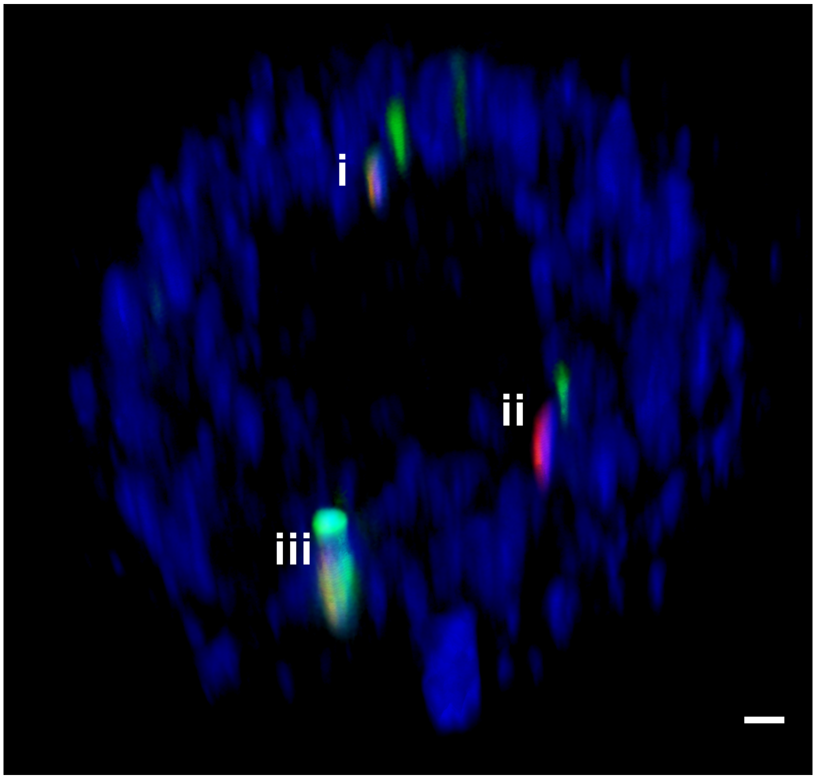


**Figure S6. Phototagging on 3D tumorsphere**. 6 MCF10A-H2B-GFP cells (green) were identified in a z-plane of the MCF10A tumorsphere mixed with a sparse amount of MCF10A-H2B-GFP cells. 3 of the 6 GFP-expressing cells were selected for phototagging and successfully phototagged, shown in red (i-iii). The whole tumorsphere was stained with Deep Red NuclearMask stain (shown in blue). Scale bar 25 µm.

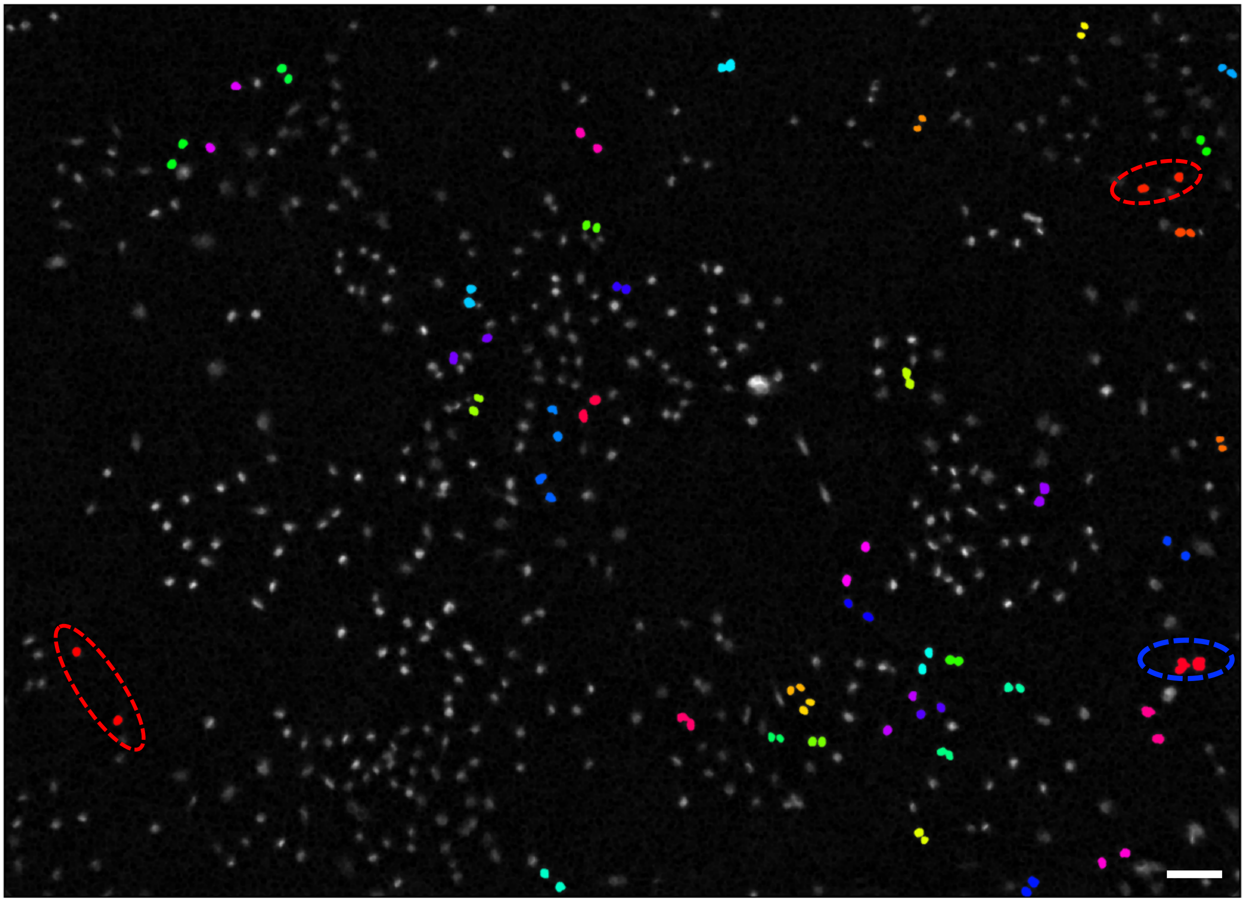

**Figure S7. Detection of cell division.** 41 pairs of newly dividing cells were in this filed-of-view. 40 were successfully detected via mTGMM and 1 pair was missed (blue dash circle). In addition, 2 pairs were falsely detected (red dash circle). Scale bar 100 µm.

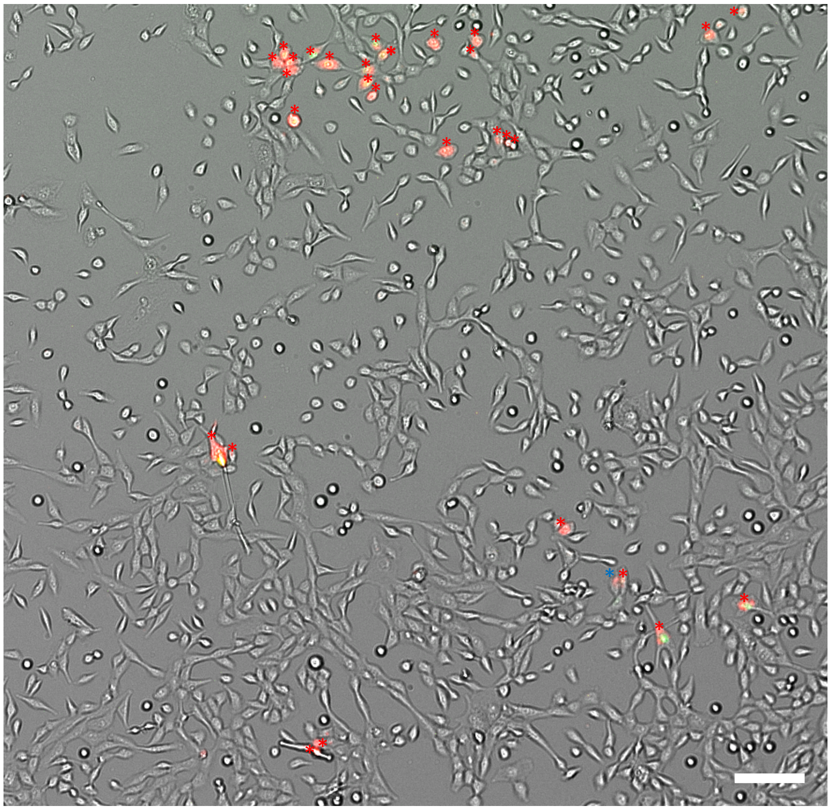

**Figure S8. Phototagging specificity**. A sparse amount (~2%) of MCF10A-H2B-GFP cells were co-cultured with MCF10A cells without GFP expression. In this representative image, 29 cells expressing GFP were successfully phototagged (red, also highlighted with *) and 1 cell was falsely phototagged (blue *). Scale bar 100 µm.

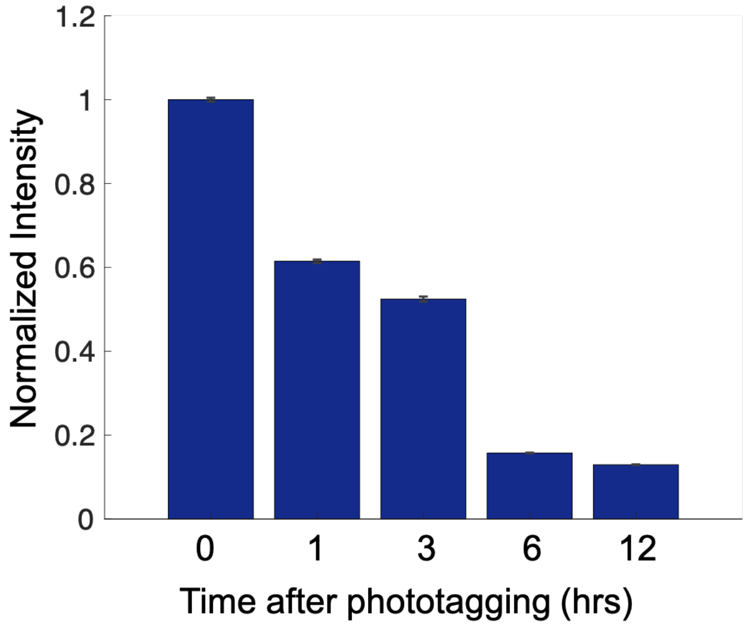


**Figure S9. Phototagging dye retention time**. Normalized intensity of phototagging dye at different timepoints after phototagging. Error bars indicate s.e.m. (N=200 cells).

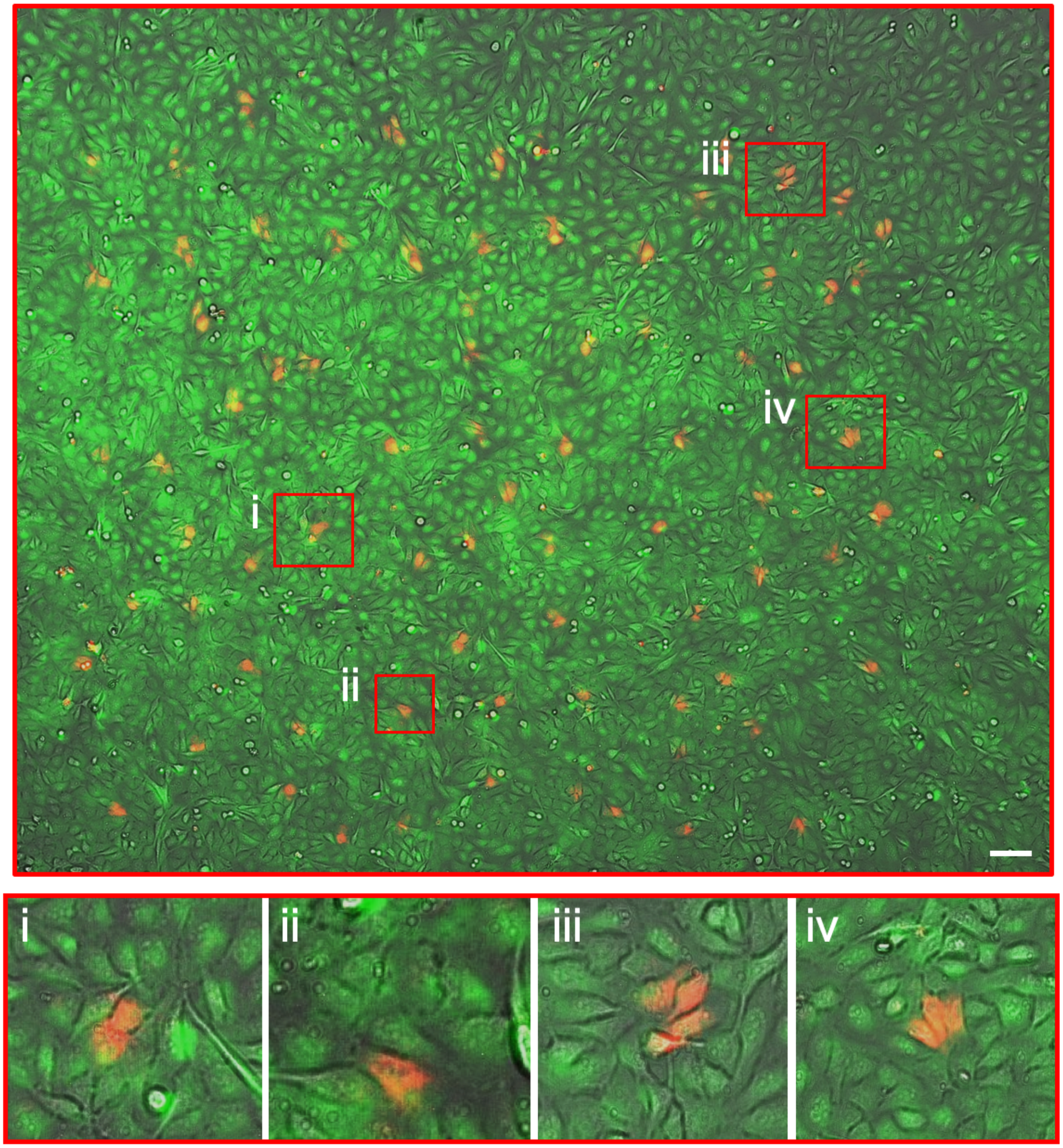


**Figure S10. MCF10A cells remain viable after phototagging**. 140 random cells were phototagged (red). After the phototagging procedure, Live Cell dye was used to stain the live cells (shown in green) in the dish. All phototagged cells (red) remain alive (green) after the procedure, as shown in the zoomed-in representative images (i-iv). Scale bar 100 µm.

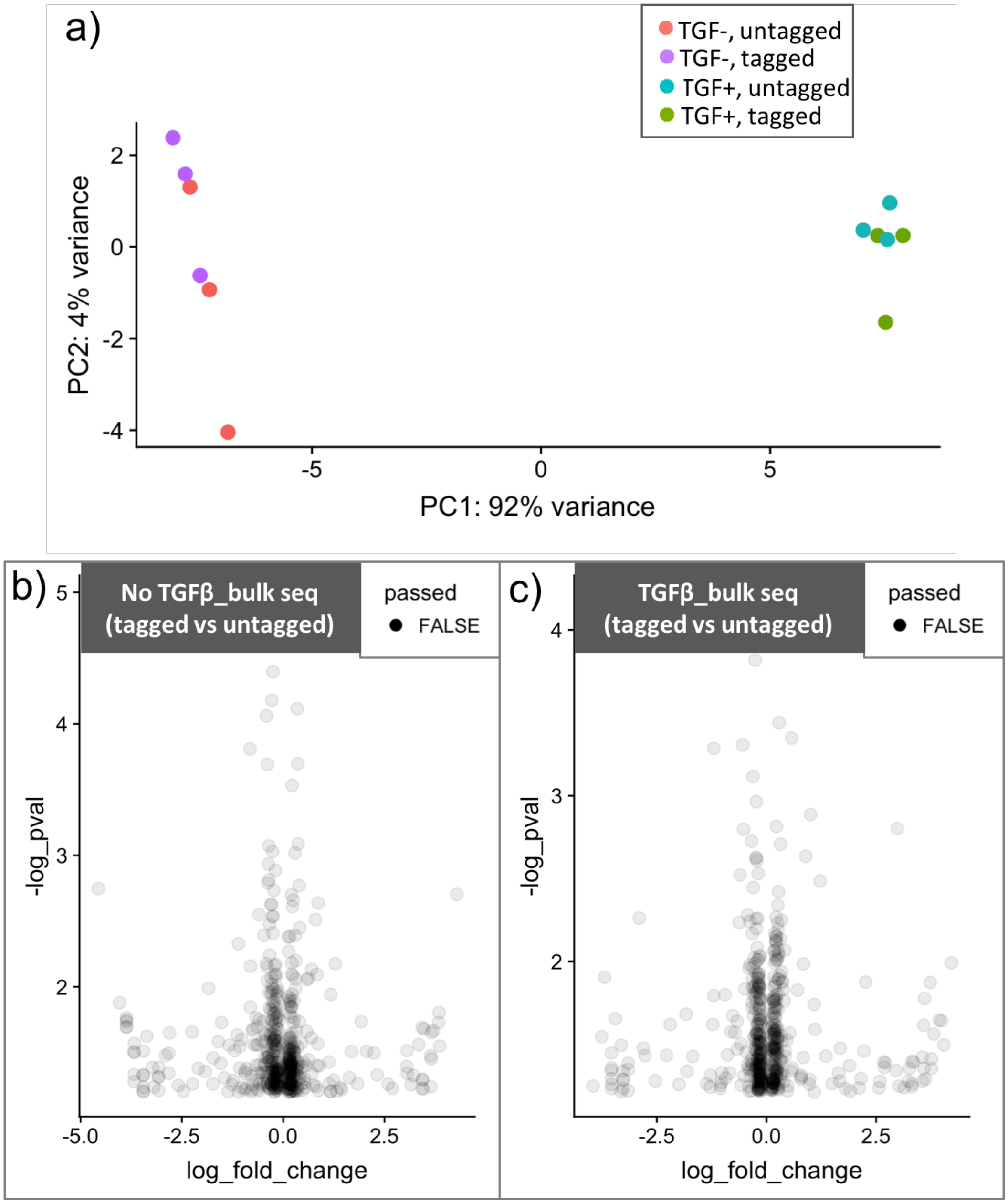


**Figure S11. PCA clustering and Volcano plots displaying differentially expressed genes between randomly selected tagged and untagged cells from bulk-cell RNA sequencing data of MCF10A cells induced and uninduced by TGFβ.** a) Three repeats of both conditions with randomly phototagged and non-phototagged cells were performed. Red: Non-TGFβ-induced, non-phototagged cells (untagged). Purple: Non-TGFβ-induced, phototagged cells (tagged). Blue: TGFβ-induced, non-phototagged cells (untagged). Green: TGFβ-induced, phototagged cells (tagged). b) Volcano plot of differentially expressed genes between randomly tagged and untagged cells from the uninduced MCF10A cells. No gene passed adjusted p-value testing (Bonferroni adjusted P value < 0.05). c) Volcano plot of differentially expressed genes between randomly tagged and untagged cells from the TGFβ-induced MCF10A cells. No gene passed adjusted p-value testing (Bonferroni adjusted P value < 0.05).

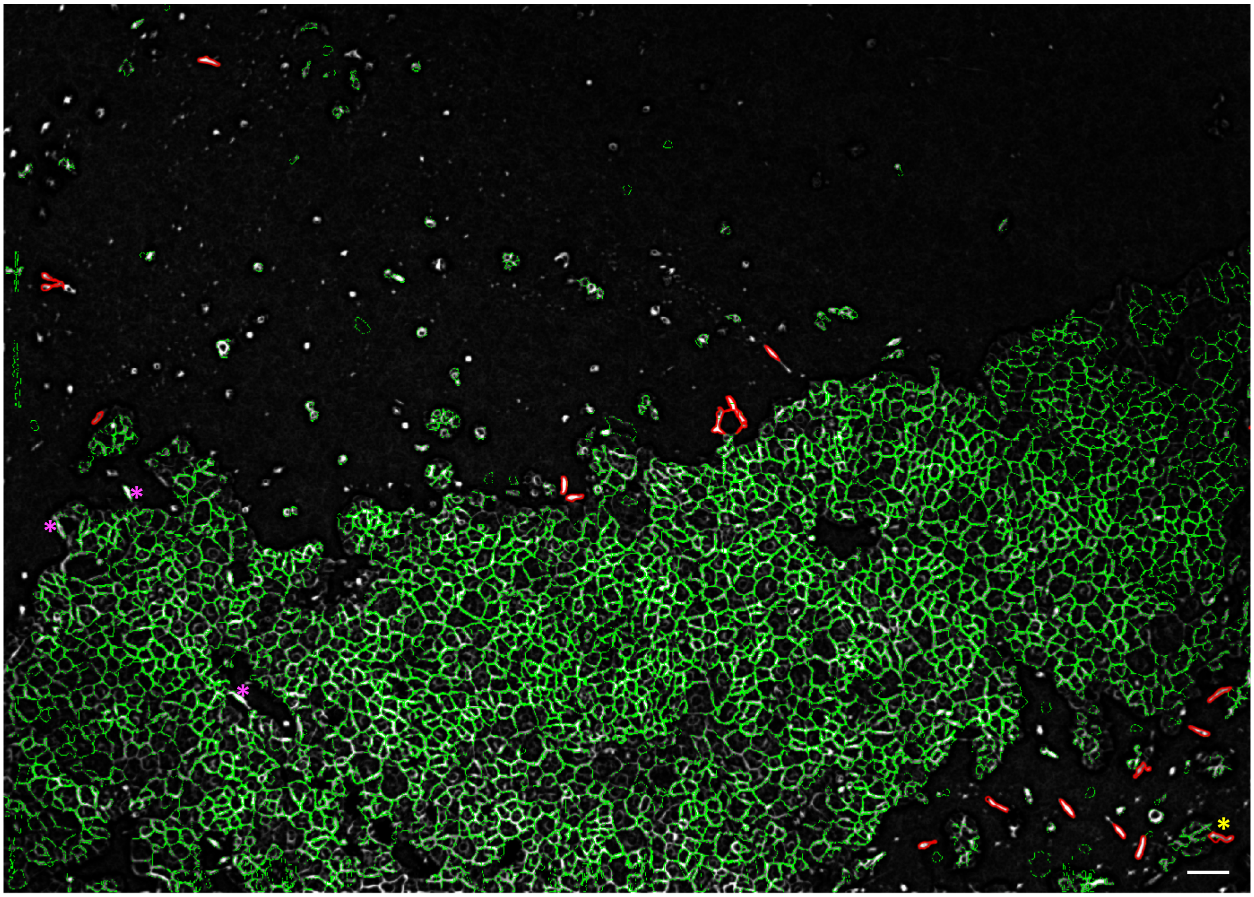

**Figure S12. Detection of cells with mesenchymal-like morphology.** 22 of spindle cells were in this field-of-view. 19 were successfully detected via mTGMM and 3 cells were missed (magenta*). In addition, 1 cell was falsely detected (yellow*). Scale bar 200 μm.

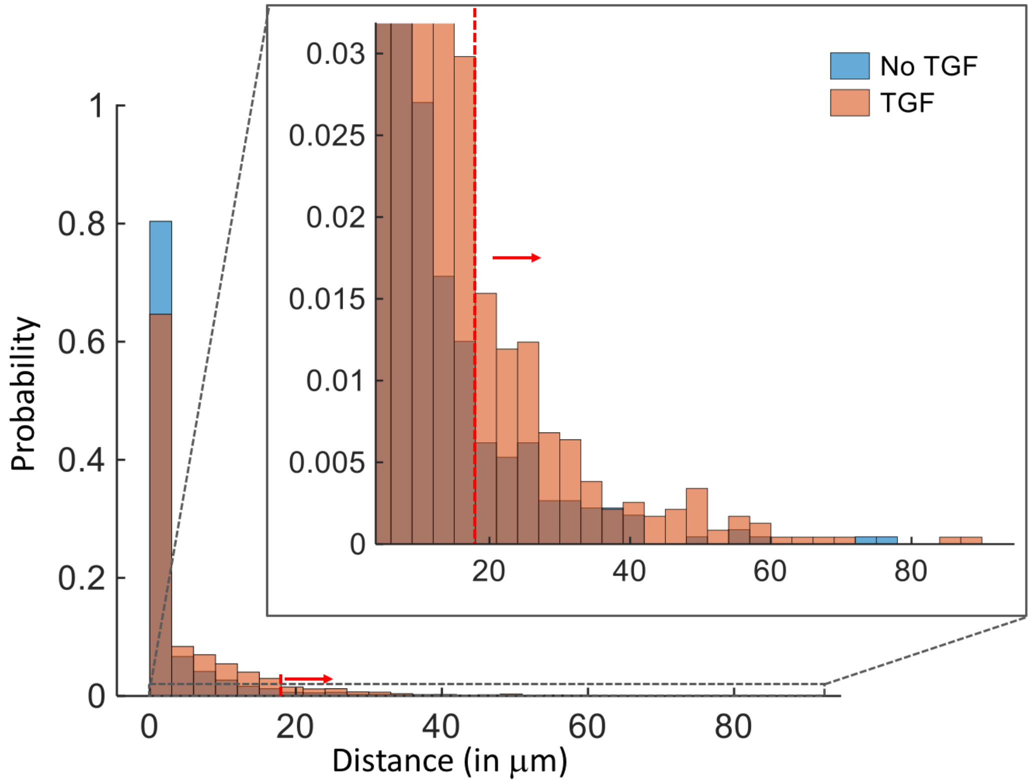


**Figure S13**. **Histogram of migration distance of TGFβ induced and non-induced cells**. Red arrow indicates the threshold (10 pixels (1.7 μm/pixel), 17 μm) used to separate faster migrating cells (after an hour of movement).

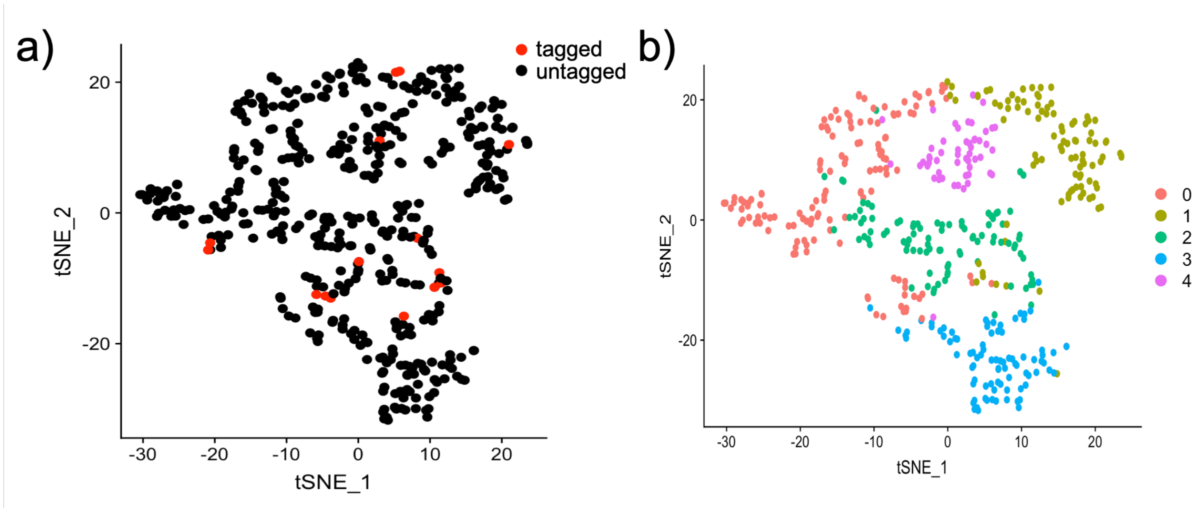


**Figure S14. tSNE plots of single cell transcriptomic profile of MCF10A-H2B-GFP cells shows no cluster based on sparse fast migrating cells in unenriched samples. a)** tSNE plot of 3% tagged cells (fast cell, red) in a large pool of slow cells (97%, black). **b)** Number 0-4 indicate the cluster number IDs identifid via Seurat's SNN-clustering method (the cell distribution matches the tSNE plot shown in figure a).

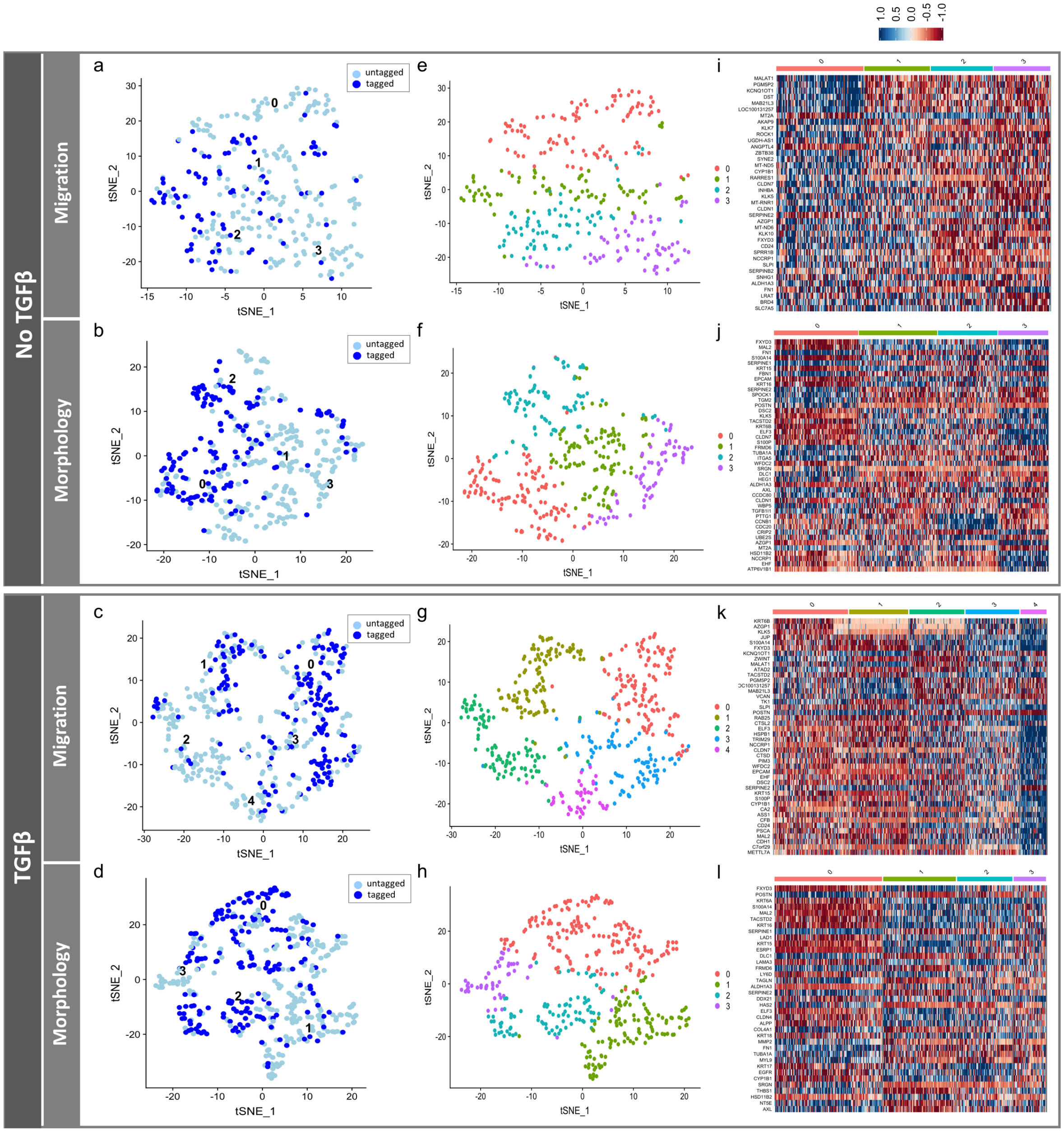


**Figure S15. Functionally annotated single cell transcriptomic profiling of MCF10A-H2B-GFP cells with mesenchymal-like morphology and fast migration with cell cycle correction. a-b, e-f, i-j): non-TGFβ induced data: a, b)** tSNE plots of non-TGFβ induced fast migrating (a) and spindle (b) cells. Dark blue: cells with fast migration or mesenchymal-like morphology (tagged); light blue: cells with slow migration or without mesenchymal-like morphology (untagged). Number 0-3 (a,b) indicate the cluster number IDs identifid via Seurat's SNN-clustering method shown in figure **e** and **f**. **i, j)** Heatmap plots of differentially expressed genes of all clusters identified by SNN for non-TGFβ induced fast migrating (i) and spindle (j) cells. **c-d, g-h, k-l): TGFβ induced data**: **c, d)** tSNE plots of TGFβ induced fast migrating (c) and spindle (d) cells. Dark blue: cells with fast migration or mesenchymal-like morphology (tagged); light blue: cells with slow migration or without mesenchymal-like morphology (untagged). Number 0-4 (c) or 0-3 (d) indicate the cluster number IDs identifid via Seurat's SNN-clustering method shown in figure **g** and **h**. **k, l)** Heatmap plots of differentially expressed genes of all clusters identified by SNN for TGFβ induced fast migrating (k) and spindle (l) cells. Differentially expressed genes between different clusters identified from SNN clustering were computed through *FindAllMarkers* function using MAST (Bonferroni adjusted P value < 0.05).

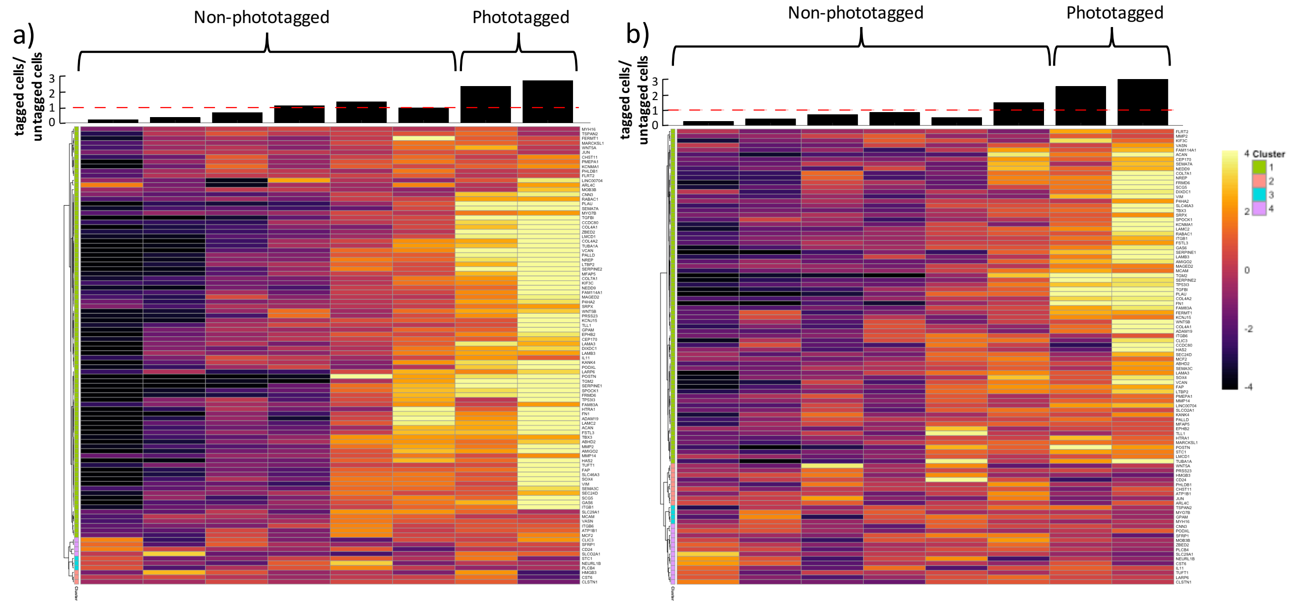


**Figure S16. Supervised EMT analysis to assess the coherence of the EMT gene module in single cell data (migration data)**. Single cells were stratified into eight clusters by EMT score, gene expression of each gene in each cell cluster was estimated and the gene trend was grouped accordingly. The proportion of tagged (fast) cells versus untagged (slow) cells is displayed on top of each column. a) TGFβ-induced data; b) Non-TGFβ induced data. Phototagged cells cluster at higher EMT scores.

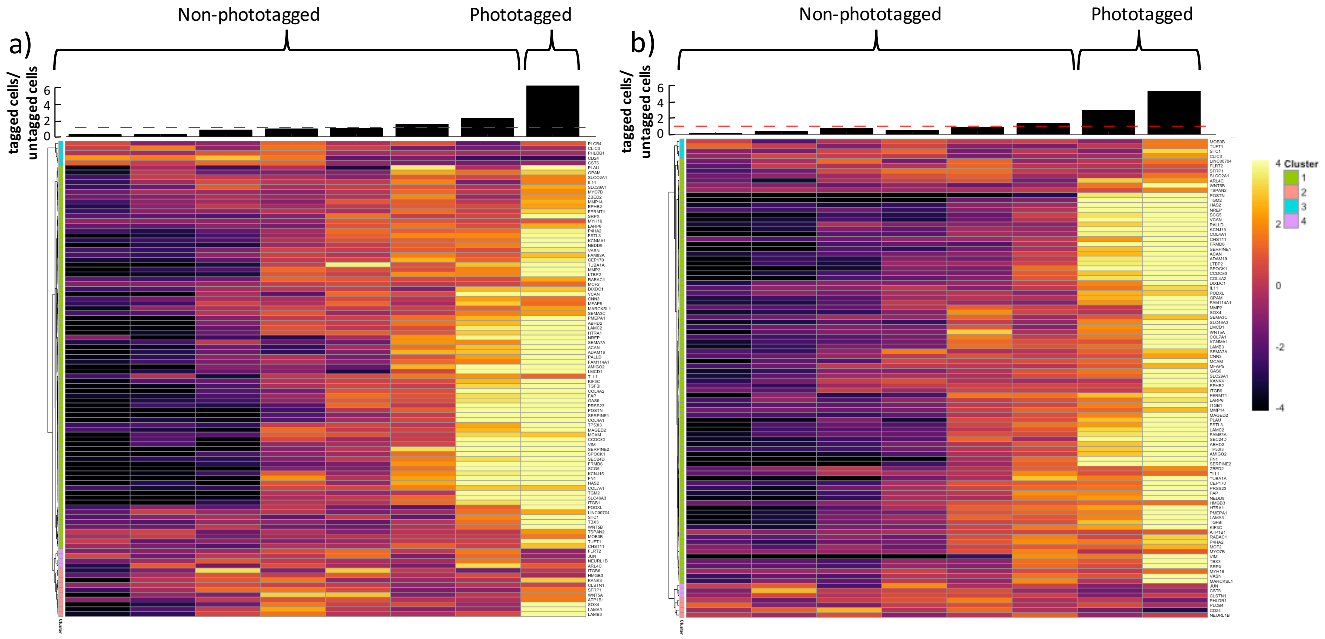


**Figure S17. Supervised EMT analysis to access the coherence of the EMT gene module in single cell data (morphology data)**. Single cells were stratified into eight clusters by EMT score, gene expression of each gene in each cell cluster was estimated and the gene trend was grouped accordingly. The proportion of tagged (spindle) cells versus untagged (non-spindle) cells is displayed on top of each column. a) TGFβ-induced data; b) Non-TGFβ induced data. Phototagged cells cluster at higher EMT scores.

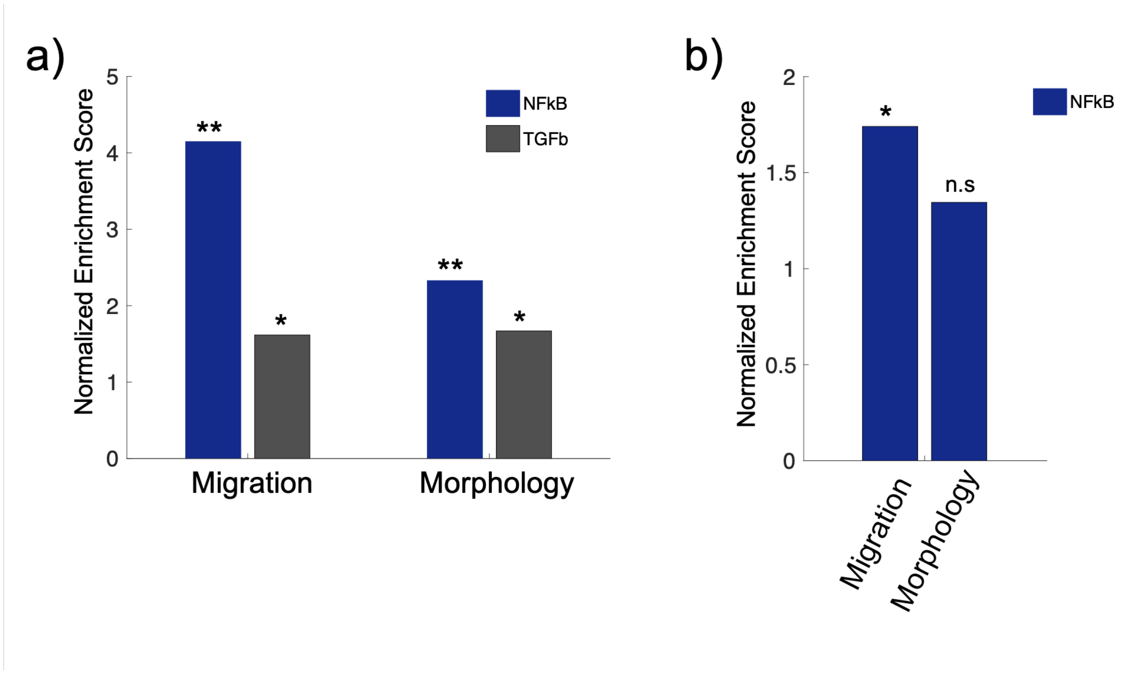


**Figure 18. Bar plots of Hallmark pathway GSEA results for the differentially expressed genes identified by comparing tagged to untagged cells**. The y-axis represents the normalized enrichment score of the gene sets associated with either NF-κB or TGFβ pathway in non-TGFβ induced MCF10A cells **(a)** or TGFβ induced MCF10A cells **(b)**. X-axis indicates the dataset of either migration or morphology phenotype. (**: passed Bonferroni adjusted P value < 0.01; *: passed Bonferroni adjusted P value < 0.05)

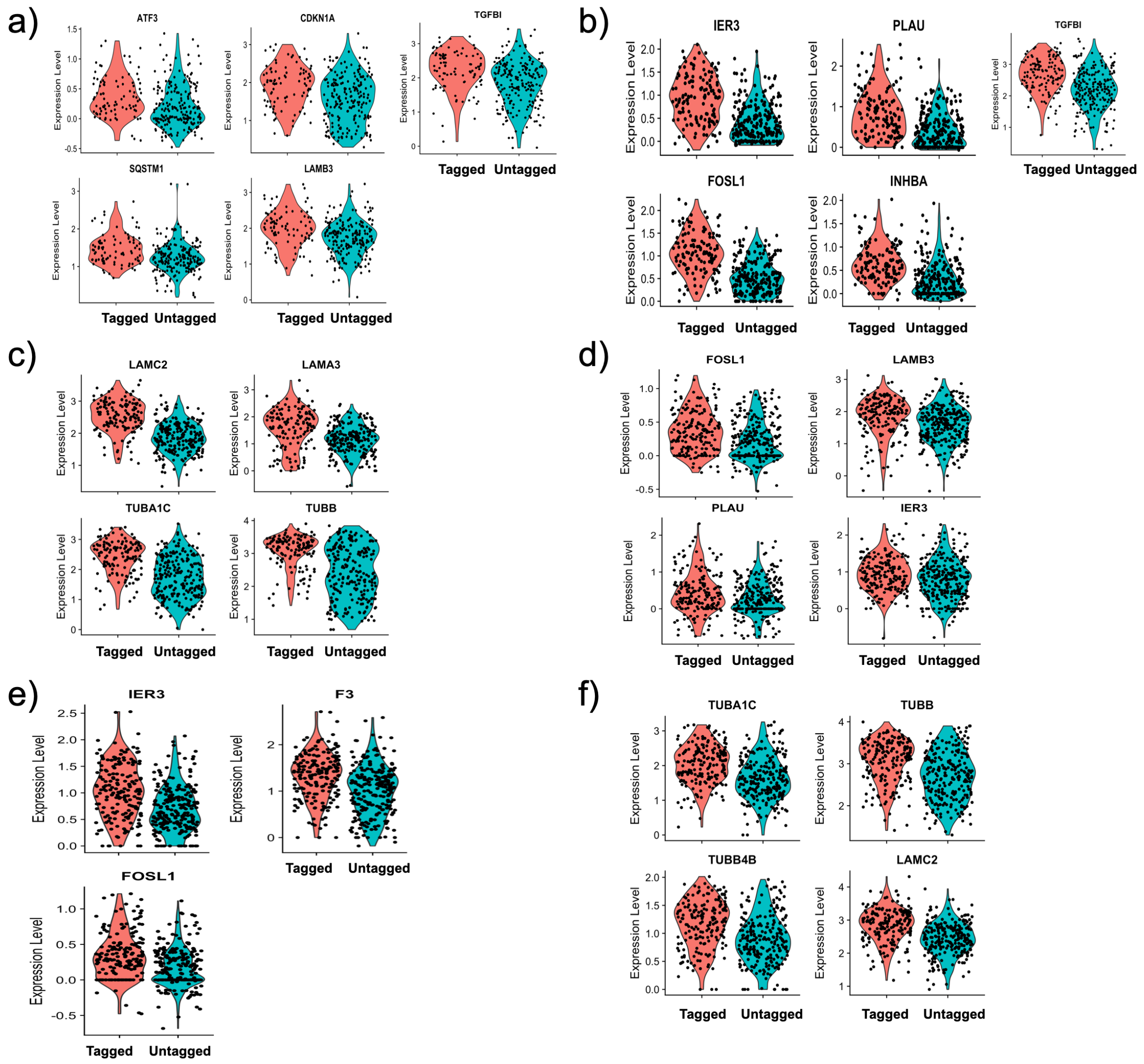


**Figure S19. Violin plots of representative differentially expressed genes identified by comparing tagged (fast or spindle) to untagged (slow or non-spindle) cells**. **a)** Differentially expressed genes between fast and slow migrating MCF10A cells in the non-TGFβ induced condition. **b)** Differentially expressed genes between spindle and non-spindle MCF10A cells in the non-TGFβ induced condition. **c)** Differentially expressed genes related to structural proteins between spindle and non-spindle MCF10A cells in the non-TGFβ induced condition. **d)** Differentially expressed genes between fast and slow migrating MCF10A cells in the TGFβ induced condition. **e)** Differentially expressed genes between spindle and non-spindle MCF10A cells in the TGFβ induced condition. **f)** Differentially expressed genes related to structural proteins between spindle and non-spindle MCF10A cells in the TGFβ induced condition. All shown genes in the violin plots passed adjusted p-value testing (Bonferroni adjusted P value < 0.05).

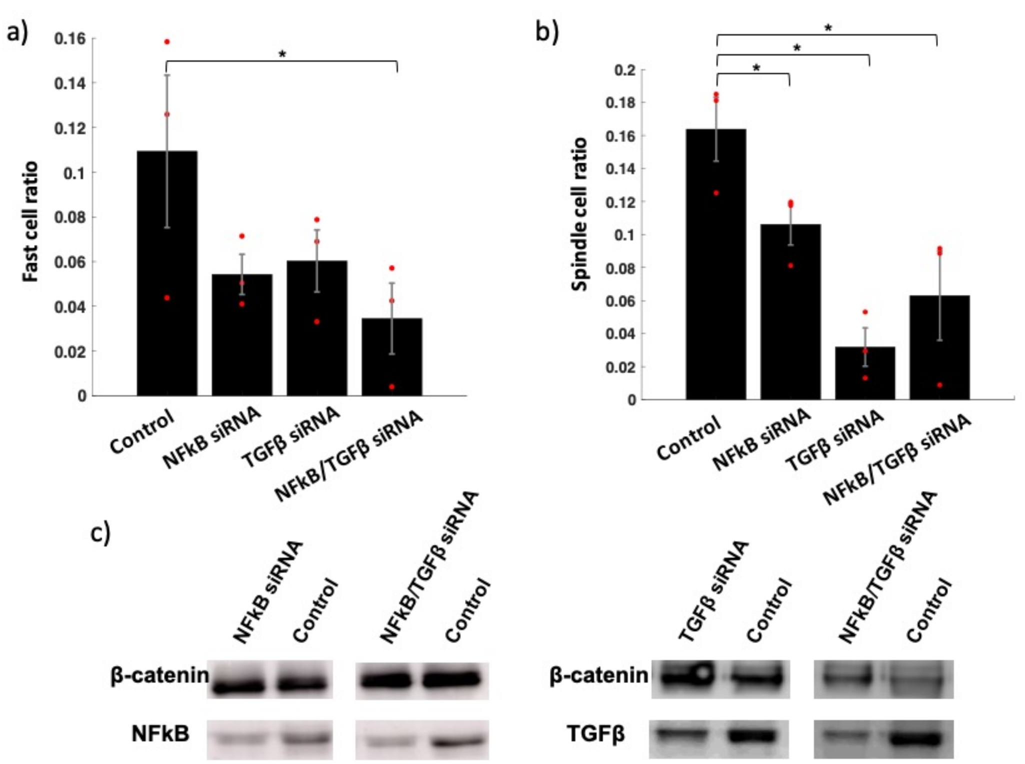


**Figure S20**. **Gene knockdown assay (RNA interference assay) of MCF10A cells treated with NF-κB and/or TGFβ siRNA. a) Migration data**: Percentage of fast migrating cells (> 10 pixels/hr; 17 μm/hour) in MCF10A cells treated with scrambled siRNA (Control), NF-κB siRNA, TGFβ siRNA or a combination of NF-κB and TGFβ siRNA. Significance (*) was computed by the ANOVA test. In comparison to the control group (scrambled siRNA), the population of fast migrating cells (17 μm (10 pixels)/hr) decreased 45% and 50% in the TGFβ and NF-κB siRNA-treated groups, respectively, and decreased 68% (with significance) in the group treated with both TGFβ and NF-κB siRNA (**a**). **b) Morphology data**: Percentage of spindle cells (mesenchymal-like morphology) in MCF10A cells treated with scrambled siRNA (Control), NF-κB siRNA, TGFβ siRNA or a combination of NF-κB and TGFβ siRNA. Significance (*) was computed by ANOVA test. **c)** Western blot of NF-κB and TGFβ proteins after the gene knockdown with scrambled (Control), NF-κB and/or TGFβ siRNA. β-catenin was used as the internal control protein. In comparison to the control group (scrambled siRNA), a decrease of spindle-shaped cells was observed of 81% in the group treated with TGFβ siRNA, 35% in the NF-κB siRNA treated group and 62% for the group treated with both siRNA (with significance) (**b**).

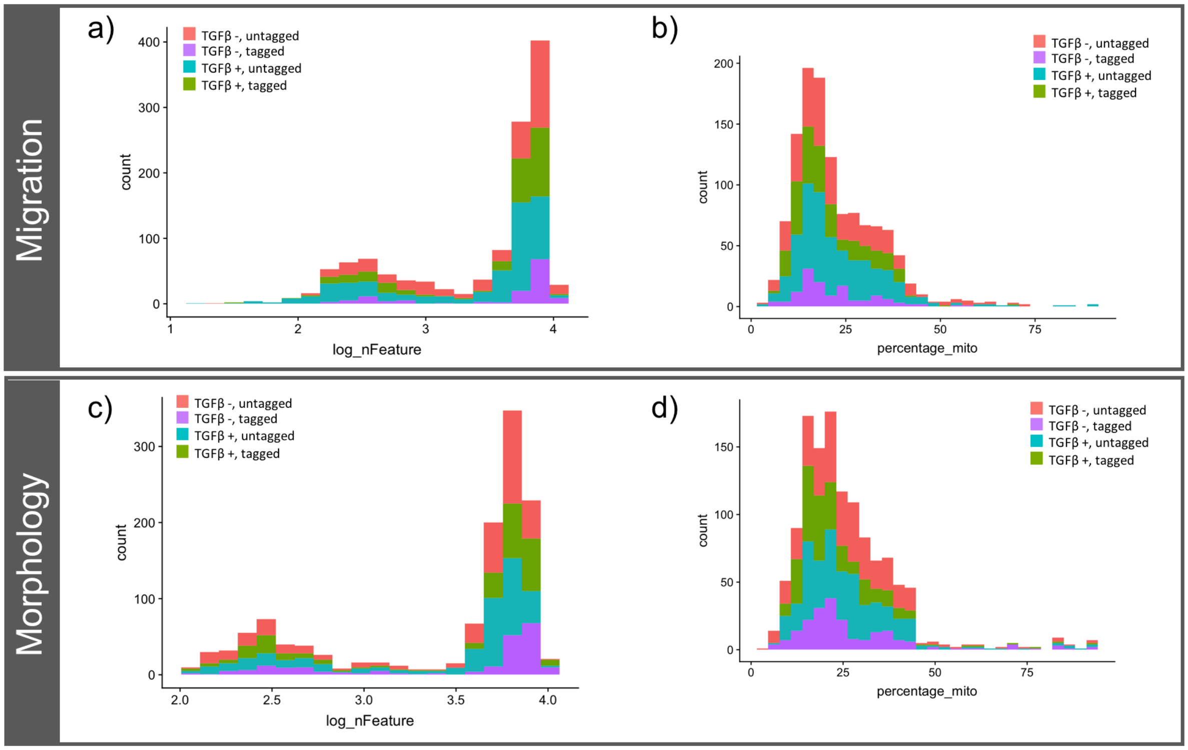


**Figure S21**. **Histograms of quality control metrics for single cell sequencing data.** a, b) Migration data: a) Log of the total feature counts. b) Percentage of mitochondrial counts. c, d) Morphology data: c) Log of the total feature counts. d) Percentage of mitochondrial counts.

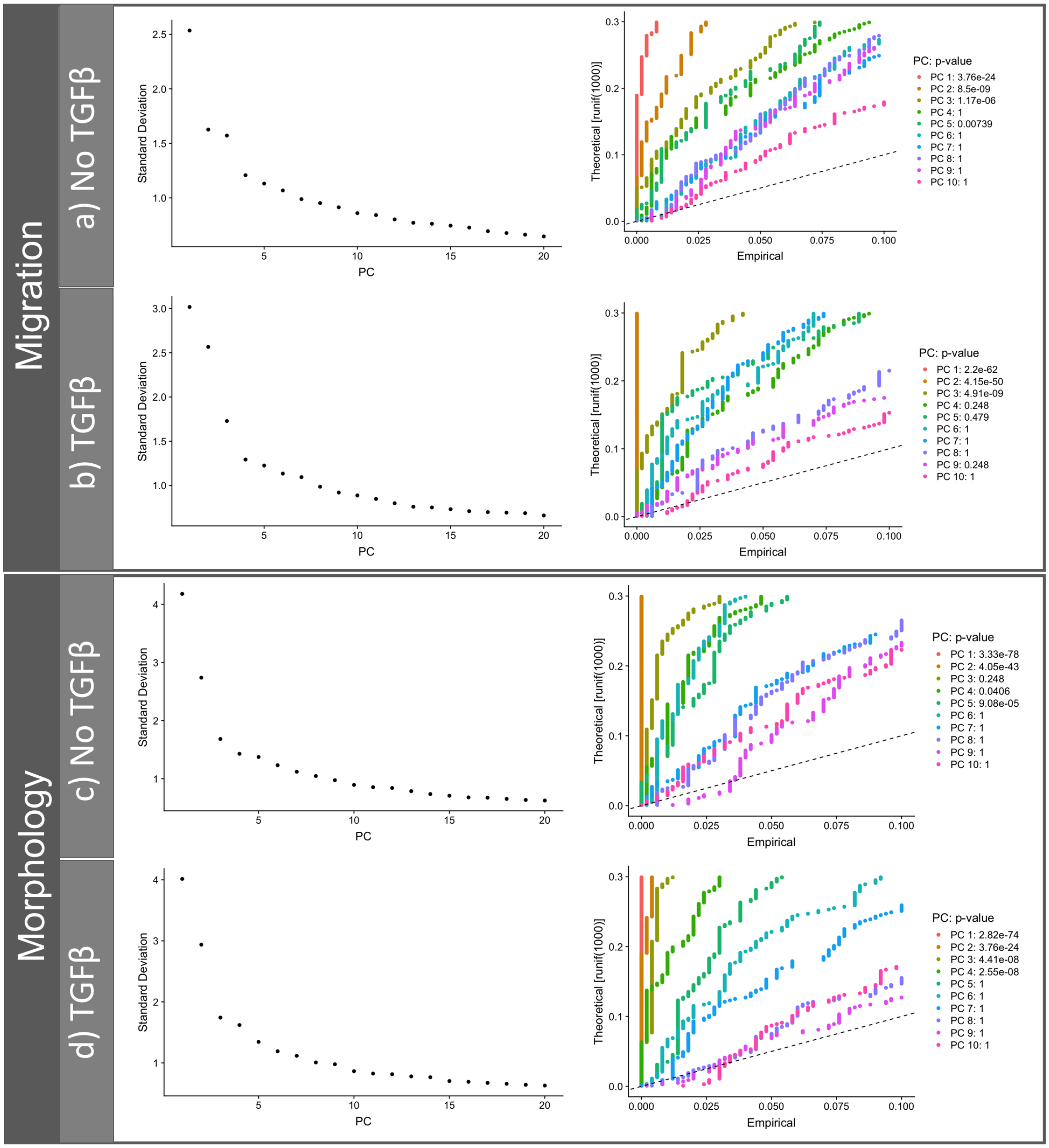


**Figure S22. Elbow plot (left) and JackStraw (right) plot to determine the amount of principle components in SNN-clustering.** a, b) Migration data: a) Non-TGFβ induced samples. b) TGFβ induced samples. c, d) Morphology data: c) Non-TGFβ induced samples. d) TGFβ induced samples. Jackstraw Plots show the cumulative distributions of p-values associated with enrichment of a gene in the principal component. Relevant principal components should be enriched for low p-values.

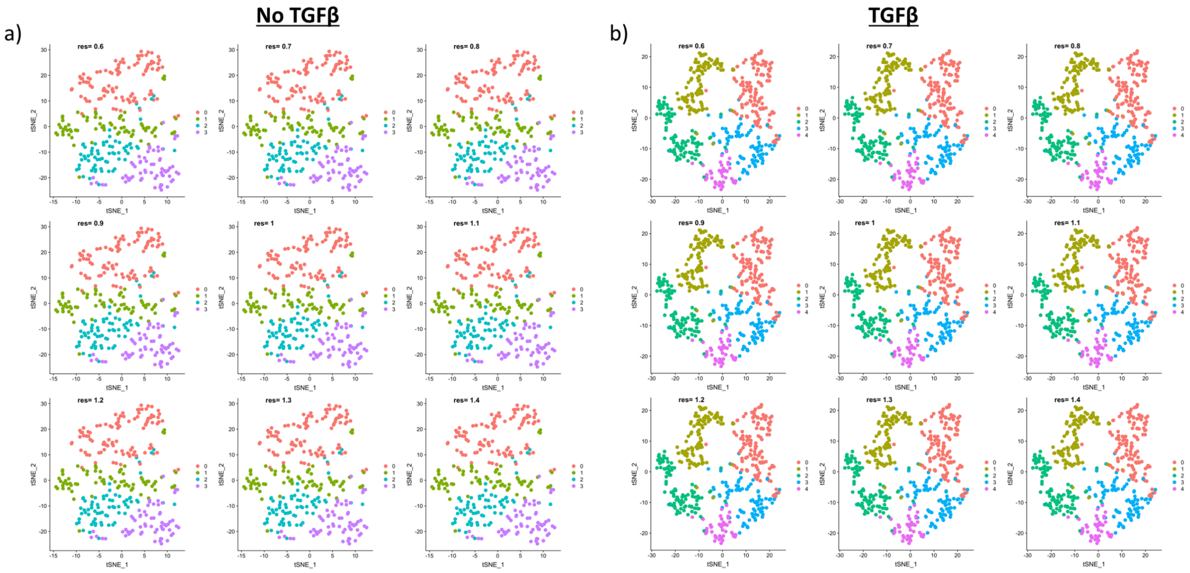


**Figure S23.** SNN-clustering of non-TGFβ induced (a) and TGFβ induced (b) cells (*migration data*). The resolution parameter was swept in the range of 0.6 to 1.4.

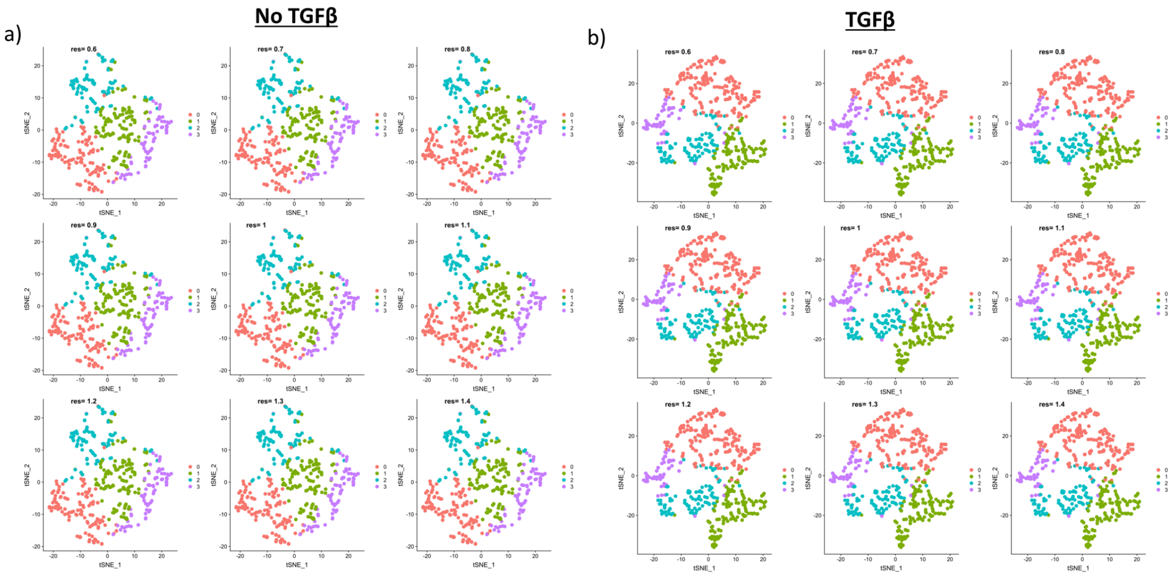


**Figure S24.** SNN-clustering of non-TGFβ induced (a) and TGFβ induced (b) cells (*morphology data*). The resolution parameter was swept in the range of 0.6 to 1.4.

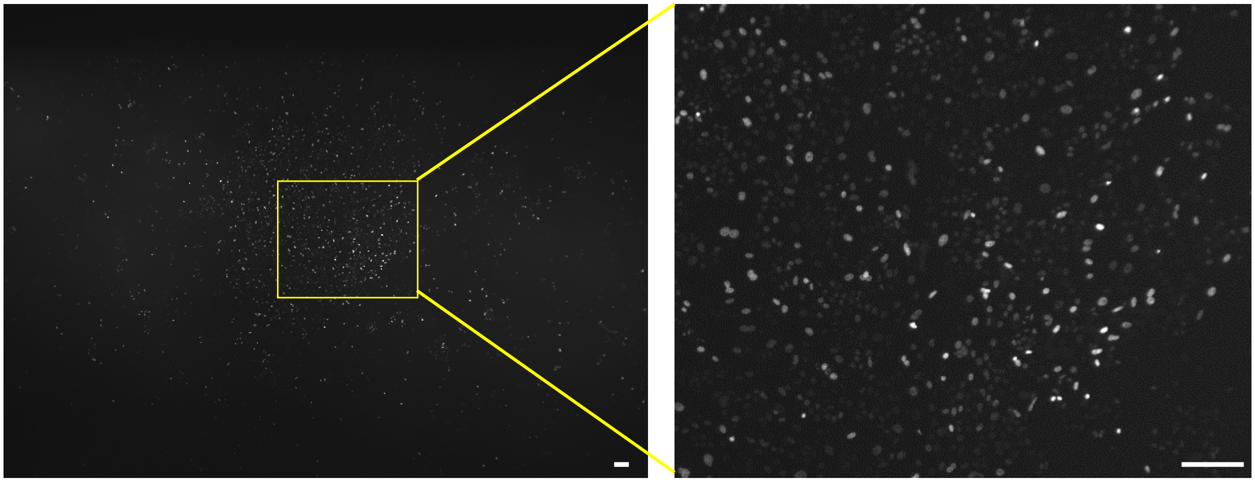

**Figure S25. A typical 2D sample where heterogeneity in fluorescence intensity is normally seen.** Left: a full field of view image from MCF10A-H2B-GFP cells. Right: Zoomed -in image of the highlighted yellow area shown in the left side. Scale bar 100 µm.

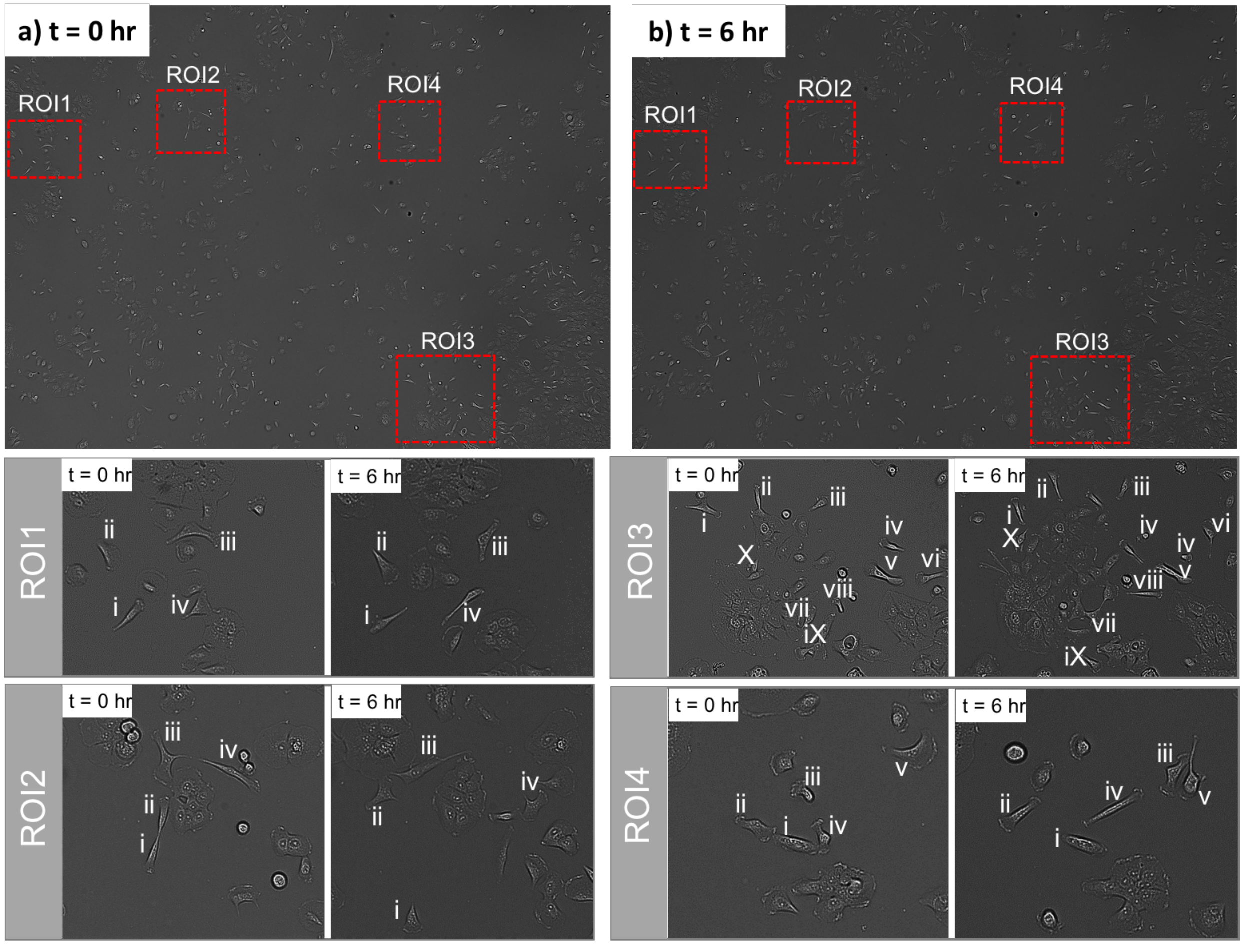


**Figure S26. Cellular morphology of MCF10A cells.** a) A full-of-view of bright-field image of MCF10A cells at t = 0 hr. b) A full-of-view bright-field image of MCF10A cells at t = 6 hr. ROI (region-of-interest) 1-4: Four zoomed-in representative ROIs from a) and b) are indicated. Most of the cells with mesenchymal morphology maintain the spindle shape after 6 hr.
